# Bridging Simplicity and Depth in Single-Cell Proteomics: A Cost-Effective Workflow and Expanded Framework for Data Evaluation

**DOI:** 10.64898/2026.02.08.700933

**Authors:** Shuxin Chi, Jason C. Rogalski, Huan Zhong, Esperanza Garcia, Arpa Ebrahimi, Rachel Wong, Melanie L. Bailey, Marco A. Marra, Claudia S. Maier, Terrance P. Snutch, Leonard J. Foster

## Abstract

Single-cell proteomics (SCP) offers direct insight into functional protein states that drive cellular heterogeneity, complementing genomic and transcriptomic analyses. Although recent reports have demonstrated improved proteome coverage, their reliance on specialized instrumentation limits broader adoption. Additionally, current evaluation practices remain largely centered on protein and peptide identification counts, which alone do not fully reflect data quality or biological interpretability. Here, we describe an accessible, label-free SCP workflow which implements easily accessible laboratory equipment: a single-cell dispenser, conventional multiwell plates, and an incubator with water-bath-based humidity control. Using trapped ion mobility spectrometry-time-of-flight mass spectrometry (timsTOF), we systematically optimize key sample preparation variables, including trypsin concentration, incubation time, reduction/alkylation, digestion conditions, and plate types, which together maximize data quality and reproducibility. We further introduce a data-quality framework that moves beyond identification counts, emphasizing quantitative consistency and biological interpretability via individual protein coverage completeness across cells, coefficients of variation across technical replicates, peptide-to-protein ratios, and single-cell-to-bulk correlations. Collectively, our approach lowers technical barriers to accessing SCP while enabling more rigorous, interpretable, and scalable SCP analysis across diverse research contexts.

## Introduction

Advancements in single-cell genomics and transcriptomics over the past decade have revealed insights into biological complexity, highlighting cellular and molecular heterogeneity as a key driver of tissue function and personalized therapeutic treatments^1–3^. Despite these breakthroughs, genomic and transcriptomic data alone cannot provide a full picture of cellular heterogeneity since proteins play crucial roles in mediating cellular functions and phenotypes. To address this gap, single-cell proteomics (SCP) has emerged as an essential complementary approach, enabling direct measurement of protein repertoires at the single-cell level.

Recent SCP methods have achieved significant milestones, particularly through the development of single-cell-specific technologies, enabling cell isolation, lysis, and protein digestion in a temperature and humidity-controlled manner^4–6^. Other advanced tools allowing low volume sample preparation using microfluidic sorters and miniaturized devices have also emerged, such as nanoPOTs^7,8^ (Nanodroplet Processing in One pot for Trace Samples). These tools demonstrate the potential for increasing proteome coverage in single cells, pushing SCP closer to widespread applications in biomedical research. Despite such technological successes, broader adoption of SCP remains challenging due to the requirement for costly reagents, or specialized instruments, limiting the broader impact of SCP. Furthermore, while studies have explored cost-effective instrumentations for SCP implementation, there remains no consensus sample preparation workflow^9,10^. Critical preparation steps, such as reduction and alkylation, trypsin/protein ratios, incubation times, and various other digestion conditions, are inconsistent across studies, illustrating the need for additional research^11–14^.

A further notable limitation of SCP implementation is how data quality is assessed. Conventional reliance on protein and peptide identification counts provides only a partial view of dataset quality, often overlooking both interpretability and experimental robustness. Other data quality assessment parameters that evaluate reproducibility, reliability, biological relevance and quantitative accuracy need to be considered.

To address these challenges, we first focus on optimizing robust and effective SCP methods accessible to general laboratory settings using trapped ion mobility spectrometry (tims) and time-of-flight (TOF) mass spectrometry in a label-free approach. Our main aim is to establish a reliable and reproducible workflow which requires minimal in-house engineering and sample preparation equipment, to improve accessibility and data quality. We further demonstrate how sample preparation procedures can be modified to address different biological questions. Lastly, we propose a comprehensive framework for evaluating SCP data which includes measures of reproducibility, reliability, and quantitative depth. Here, we refer to reproducibility as the consistency of protein detection and quantification across single cells, evaluated through the coefficient of variation (CV) across replicates and individual protein coverage completeness; reliability by the confidence in protein assessments as measured by the peptide-per-protein ratio and their corresponding median intensity; and quantitative depth as the overall scope and robustness of protein quantification, assessed by dynamic range of protein intensity, peptide length distribution and concordance with bulk proteomics^15^.

## Methods

### Cell culture

#### A549

A549 (ATCC CCL-185) cells were cultured at 37°C with 5% CO_2_ and maintained with Dulbecco’s Modified Eagle’s Medium (DMEM; GIBCO 12800-017, Thermo Fisher Scientific) containing 10% FBS (Gibco12483020), 1% penicillin/streptomycin, and 1% sodium pyruvate. The cells were subcultured every two days and harvested at 80% confluency.

#### HEK293

HEK293-F cells (Invitrogen 11625-019) were grown in DMEM supplemented with 10% heat-inactivated FBS (hi-FBS; inactivated for 30 min at 56°C) and 1% non-essential amino acids (NEAAs; GIBCO 11140-050). Cells were maintained at 37°C in a humidified incubator at 5% CO_2_. Stable HEK293-F cell lines expressing the full-length cDNA constructs of the human Ca_V_3.1 (hCa_V_3.1) and human Ca_V_3.2 (hCa_V_3.2) T-type calcium channels were also maintained in DMEM + 10% hi-FBS and 1% NEAAs, plus 25 µg/ml of Zeocin selection antibiotic (Invitrogen Cat No. R25001). These cells were grown to 80% confluence and subcultured every two days using 0.25% trypsin-EDTA (GIBCO Cat 25200-056), replacing selection media daily.

#### Primary astrocytes

Primary astrocytes (ScienCell) were cultured Astrocyte Medium (ScienCell) at 37°C in a humidified incubator with 5% CO_2_. The cells were grown to 80% confluency used within five passages to maintain viability and health.

#### PC12

PC12 cells, obtained from ATCC (ATCC, CRL-1721), were cultured in Roswell Park Memorial Institute (RPMI) medium supplemented with 5% fetal bovine serum (FBS) (Part No. F4135; Sigma-Aldrich), 10% horse serum (HS) (2 Part No. 6050088; Thermo Fisher Scientific), and 1% penicillin-streptomycin (Part No. 15140122; Thermo Fisher Scientific). Cells were differentiated into neuronal-like cells using 100 ng/mL of Nerve Growth Factor (NGF) (Sigma-Aldrich). Following seeding, the medium was replaced with OPTI-MEM (Thermo Fisher Scientific) containing 0.5% FBS to initiate neuronal differentiation. NGF-treated cells were cultured for up to 6 days, with media and NGF replaced every two days to ensure consistent and effective differentiation. Cells were collected on days 2, 4, and 6 after treatment.

The collected cells (A549, HEK293, astrocytes, and PC12) were washed three times with 1x phosphate-buffered saline (PBS) through centrifugation at 300 x g for 5 min. The washed cells were resuspended in 1x PBS and the cell density and size were measured using CountessII. The final cell density was diluted to 2×10^5^ cells/mL. Single cells were dispensed using the Uno Single Cell Dispenser (Tecan Trading AG, Switzerland)^16^ into twin.tecTM PCR 384 well-plates (Eppendorf, Hamburg, Germany).

#### Primary monocytes

Peripheral blood mononuclear cells (PBMCs) were obtained from StemCell Technologies. To enrich for CD14+ monocytes, PBMCs were stained with an antibody mastermix containing anti-CD3 (SK7), anti-CD56 (NCAM1), and anti-CD19 (H1B19) antibodies conjugated to biotin (Biolegend). Magnetic separation was then performed magnetic separation using Dynabeads (Thermo Fisher) according to manufacturer’s protocol. We then dispensed monocytes from the flow-through into twin.tecTM PCR 384 well-plates (Eppendorf, Hamburg, Germany).

### Sample preparation

Single cells (n=15/condition) were dispensed using the Tecan Uno Dispenser into twin.tec™ PCR Plates 384 well-plates (Eppendorf, Hamburg, Germany) or regular 96 well-plates (Agilent Technologies, Santa Clara, CA, USA). Plates were pre-filled with 0.5µl LC-MS grade water (J.T.Baker, Avantor, Radnor, PA, USA) or 1.5µl digestion solution mix, containing Trypsin/LysC Mix (Promega Corporation, USA) in 50mM triethylammonium bicarbonate (TEAB) with 0.03% (w/v) n-dodecyl-β-D-maltoside (DDM). One round of freeze-thaw cycle was performed to lyse the cells. For samples pre-filled with water, 1µl of digestion mixture was added. Samples were incubated at 37°C for 1 hour, 2 hours, 4 hours, 8 hours, or 16 hours. For reduction and alkylation tests, 5mM (tris(2-carboxyethyl)phosphine) (TCEP, BioShop Canada Inc.) or dithiothreitol (DTT) and 20mM CAA were added along with trypsin-LysC mixture for a 2 h incubation. Incubation was done in a cell culture incubator with <=95% humidity or a PCR thermocycler for comparison. Selected samples also went through reduction and alkylation during digestion and quenching after digestion for comparison tests.

### Liquid chromatography

Single cell samples were injected and separated on-line using the NanoElute 2 UHPLC system (Bruker Daltonics, Germany) with Aurora Series Gen3 (CSI) analytical column, (25cm x 75μm 1.6μm C18 120Å, with CSI fitting; Ion Opticks, Parkville, Victoria, Australia). The analytical column was heated to 50°C using a column toaster M (Bruker Daltonics, Germany). Solvent A consisted of 0.1% aqueous formic acid and 0.5% acetonitrile in water, and solvent B consisted of 0.1% aqueous formic acid and 0.5% water in acetonitrile. Before each run, the analytical column was conditioned with 4 column volumes of solvent A. The analysis was performed at 0.10μL/min, 0.20μL/min, 0.30 μL/min and 0.4μL/min flow rate. The samples were run with a 7.5min, 15min, 30min, and 60min gradient.

### Mass spectrometry

A Trapped Ion Mobility - TimsTOF SCP tandem mass spectrometer (Bruker Daltonics) was used for data collection. The Captive Spray ionization source was operated at 1700 V capillary voltage, 3L/min drying gas and 200°C drying temperature. During analysis, the TimsTOF SCP was operated with Parallel Accumulation-Serial Fragmentation (PASEF) scan mode for DIA acquisition. The MS spectra were collected in positive mode from m/z 100 Th to m/z 1700 Th, and from ion mobility range (1/ K0) 0.7 V*s/cm2 to 1.3 V*s/cm2. The TIMS was operated with equal ramp and accumulation time of 65ms or 100 ms (100% duty cycle). For each TIMS cycle, 11 dia-pasef scans were used, each with 3-4 steps. A total of 36 dia-pasef windows were used spanning from m/z 299.5 Th to m/z 1200.5 Th, and from ion mobility range (1/ K0) 0.7 V*s/cm^2^ to 1.3 V*s/cm^2^ with an overlap of m/z 1 Th between two neighboring windows. The collision energy was ramped linearly as a function of mobility value from 20eV at 1/K_0_ = 0.6 V·s/cm^2^ to 65eV at 1/K_0_ = 1.6 V·s/cm2.

### Data analysis

Data were collected over the course of one year and, for consistency, all single-cell data were analyzed using DIA-NN^17^ version 1.8.2 against a human FASTA database (Uniprot UP000005640) and in-house built common contaminants. Bulk libraries were generated using 10ng injections and the same method. Single-cell samples were analyzed using a spectral library and the Match-Between-Runs (MBR) feature, leveraging bulk samples for alignment. *In silico* digestion of a FASTA database was performed using trypsin/P with allowance for one miscleavage. Deep learning-based predictions of MS/MS spectra, retention times (RTs), and ion mobilities (IMs) were enabled to enhance peptide identification. Peptide length ranged between 7-30, precursor charge ranged from 2 to 4, precursor m/z ranged from 300 to 1200, and fragment ion m/z ranged from 200 to 1800. Precursor FDR was set to 1%, with 0 for settings ‘mass accuracy’, ‘MS1 accuracy’ and ‘scan window’. Settings ‘heuristic protein inference’, ‘use isotopologues’, and ‘no shared spectra’ were all enabled. ‘Gene’ was chosen for protein inference parameter along with ‘double-pass mode’ for neural network classifier. Robust LC (high precision) was used as the quantification strategy, RT-dependent mode for cross-run normalization, and smart profiling mode for library generation. Data quality control filters were implemented by removing proteins with more than 70% missingness and samples with less than 1000 protein counts for A549, HEK293 and primary astrocytes, and 500 protein counts for PBMC cells.

### Data quality evaluation

We evaluated data quality according to protein counts, protein coverage completeness, tryptic miscleavage, peptide length distribution, coefficient of variance across replicates, dynamic range of protein intensity, and peptide-per-protein ratio (Figure 2-5 panel a-d, Suppl. figure 1,2). Protein coverage completeness quantified how many proteins are consistently identified across the cohort (i.e., detected in a specified fraction of cells). Tryptic miscleavage measured number of internal K/R residues not cleaved. Peptide length distributions summarized peptide intensity as a function of peptide sequence length. The coefficient of variation (CV) was computed for protein intensities across single cells within each condition to assess reproducibility. Dynamic range was estimated from the distribution of log10-transformed protein intensities for each preparation, enabling comparison across conditions. The peptide-per-protein ratio was calculated as the mean number of unique peptides identified per protein within each condition. Pearson correlations were calculated between each single-cell condition and bulk reference samples, with log-transformed intensities shown in supplementary tables. For reduction and alkylation assessment, enrichment analysis of unique protein groups was conducted using Gene Ontology (GO) Biological Process terms to identify pathway-level effects introduced by different sample preparation conditions.

**Figure 1:**
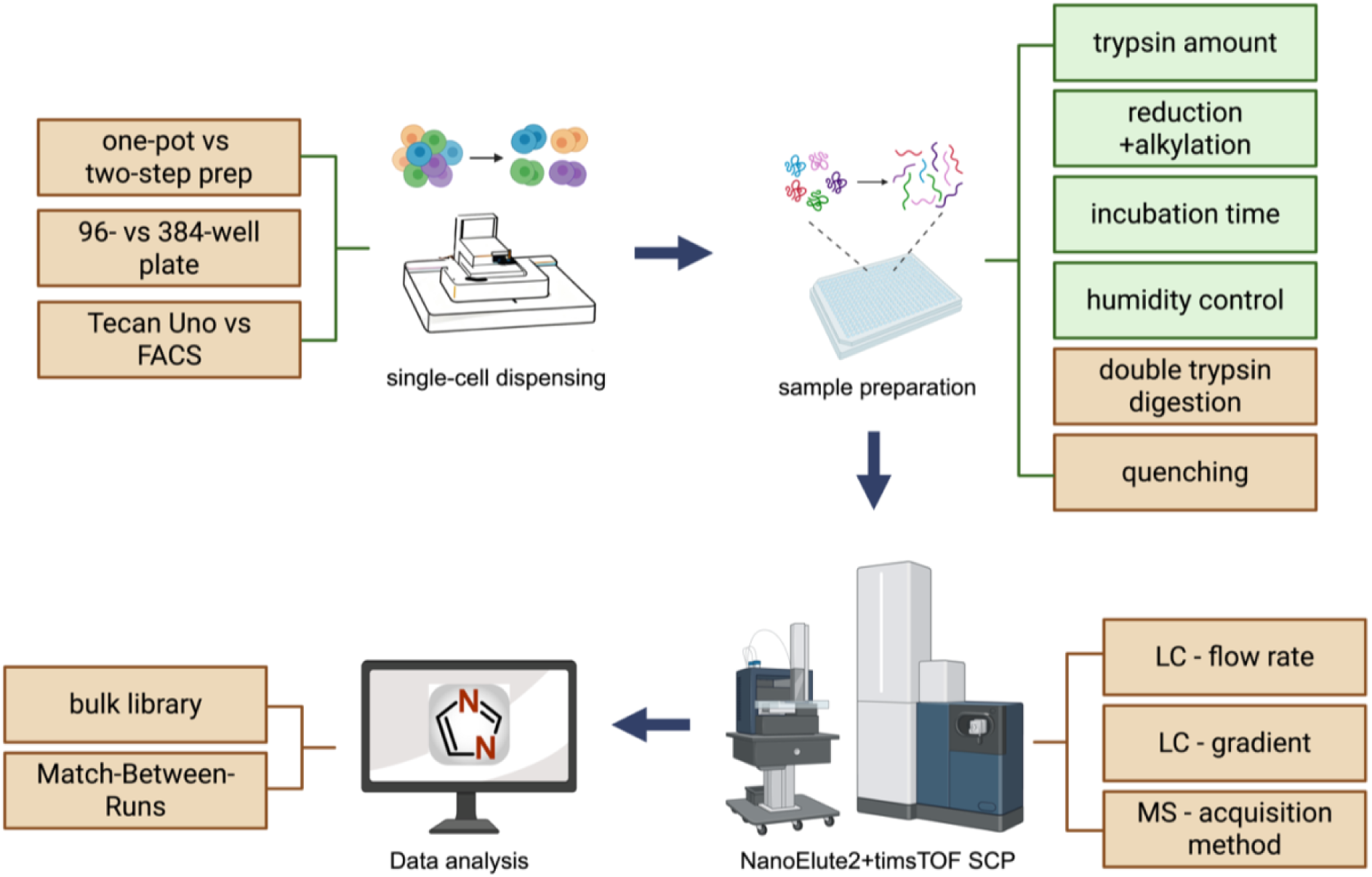
Overall workflow for single-cell proteomic optimization. The parameters tested for sample preparation are indicated in the boxes. This work focuses on the sample preparation parameters highlighted in green. (Created with BioRender.com)

**Figure 2:**
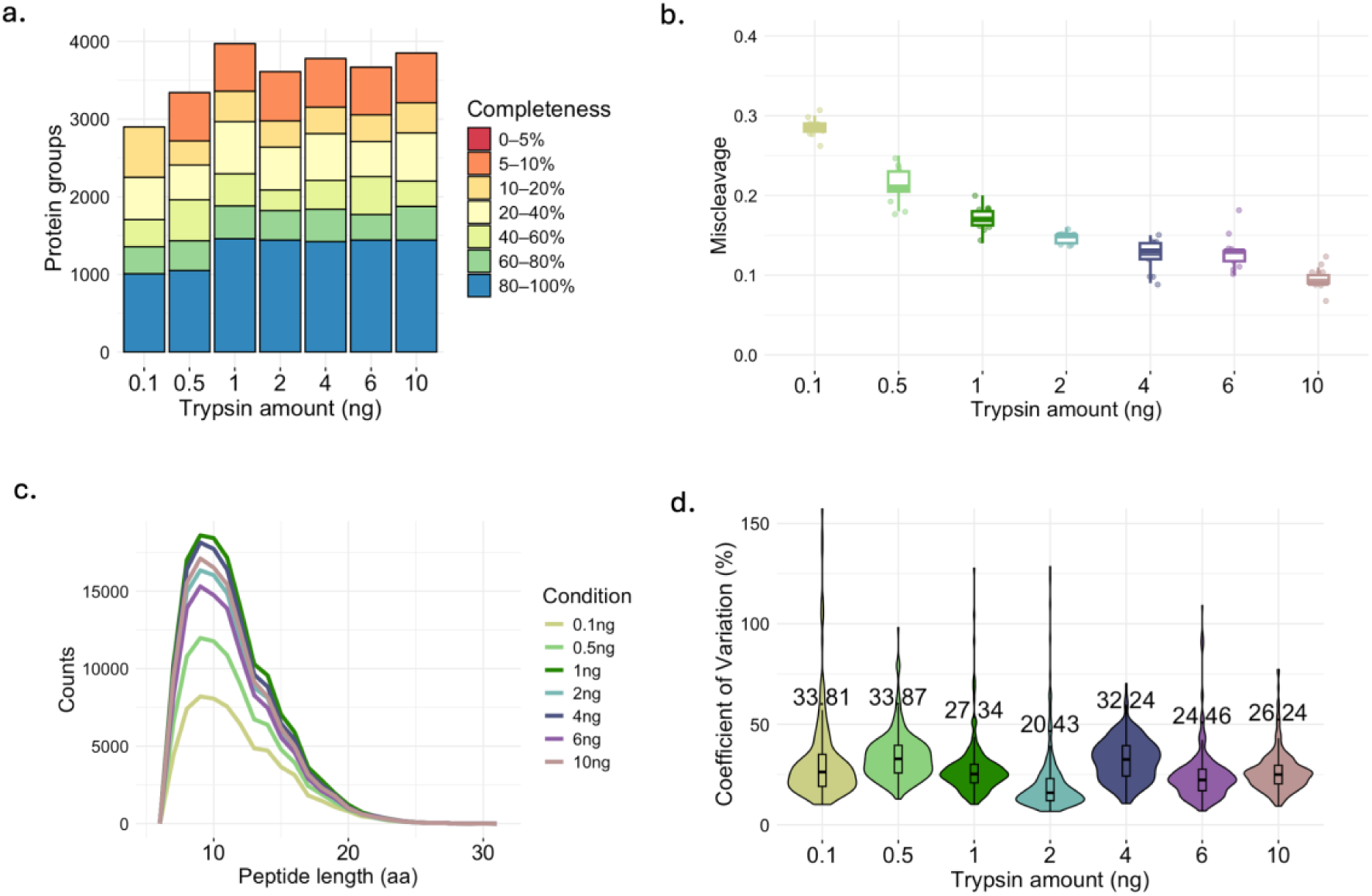
Effect of trypsin amount (0.1ng-10ng) on single-cell proteome coverage and reproducibility. a. Protein coverage completeness across cells. Stacked bars show the number of protein groups binned by completeness (0–5% to 80–100%); b. Tryptic miscleavage of each trypsin condition; c. Peptide length distributions (Peptide Spectrum Match (PSM) versus amino-acid length) across different trypsin amounts; d. Coefficient of variation (CV) of protein intensities across cells shown with violin plots with median labeled for each condition.

### Data visualization

Single-cell proteomic profiles were visualized using UMAP, computed on all quantified proteins, implemented in the scDataviz R package^18^, dot plots of tricarboxylic acid (TCA) cycle proteins, and ridgeline histograms of selected cell markers. All plots were created using R (Version 2024.09.1+394).

## Results

To design a robust and reproducible SCP sample preparation pipeline without the need of specialized equipment, we evaluated key sample preparation parameters such as trypsin amount, digestion incubation time, the inclusion/exclusion of reduction/alkylation, plate types, and digestion conditions (Figure 1). We dispensed single A549/HEK293 into a 384-well plate using a Tecan single cell dispenser. Trypsin digestion mixture was added into each sample and incubation at 37°C was conducted in a cell culture incubator with controlled humidity at <=95%. Samples were analyzed using LC-MS/MS and database searches to assess parameters including data completeness, tryptic miscleavage, peptide length distribution, coefficient of variation, protein dynamic range (Suppl. Figure 1), peptide to protein ratio (Suppl. Figure 2), and single-cell data correlation to bulk (Suppl. Table 1). Here, we focus on sample preparation parameters, LC-MS/MS optimization details were evaluated and presented in the supplementary materials.

### Single-cell sample preparation parameters compared

#### Optimizing Trypsin Load: Balancing Digestion Efficiency and Proteome Depth

The amount of trypsin for protein digestion is among the most variable parameters in single-cell proteomics workflows, with reported values covering two orders of magnitude (0.1 ng to 10 ng per cell). To evaluate the impact of the amount of trypsin on digestion efficiency for single cells ∼10-15µM in diameter, we tested 0.1 ng to 10 ng of trypsin in 384-well plates in 1µl of TEAB with 0.03% DDM using A549 with 15 cells per replicate. Our results suggest that protein coverage completeness was comparable across the 1 ng to 10 ng range (Figure 2a, Suppl. TablTable 2). Although increasing the amount of trypsin reduced trypsin miscleavages (Figure 2b), this was accompanied by a decrease in peptide detection, particularly for peptides in the range commonly detected for MS/MS analysis (7-13 amino acids)^19^ (Figure 2c). Adding an excess amount of trypsin risks limiting protein quantitative depth, possibly due to ion suppression or overloading the LC column with undigested trypsin (Suppl. Figure 3, Table 3). Other quality metrics, including coefficient of variation (CV), dynamic range of protein intensities, peptide-per-protein ratio, and correlation to bulk proteomes, were not statistically different across the range tested (Figure 2d, Suppl. Table 1, Suppl. Figure 1a,2a). While optimal enzyme load is predicted to scale with cellular protein content and may require adjustment for markedly larger or smaller cell types, for single cells ∼10-15 µm in diameter, 1 ng trypsin per well yielded high quality data without loss of proteome coverage or digestion efficiency.

### Trade-offs of Reduction and Alkylation in SCP Workflows

Reduction and alkylation are standard steps in bulk proteomics workflows that enhance protein digestion efficiency by disrupting disulfide bonds. The SCP field, however, inconsistently implemented the process during single-cell preparations. Here, we evaluated the feasibility of implementing reduction and alkylation steps in single-cell proteomics sample preparation and the efficacy of enhanced digestion on improving protein coverage completeness and the overall proteome coverage, especially for those proteins with specific disulfide bonds. Using A549 cells (n=15/condition), we compared multiple conditions: no reduction/alkylation (0 mM TCEP + CAA), full reduction and alkylation (5 mM TCEP + 20 mM CAA), and reduction-only protocols using either 5 mM TCEP or 5 mM DTT. Contrary to observations from bulk proteomics workflows, full reduction and alkylation in the context of single-cell proteomics was associated with decreased reproducibility as indicated by an approximately 50% reduction in proteins reaching 80–100% completeness and by more than 500 proteins being uniquely identified in the control (0mM TCEP+CAA) condition (Figure 3a, Suppl. Table 2). Furthermore, reduction and alkylation introduced contaminants when no pre-loading cleanup was performed (Figure 3a, 3d, Suppl. Figure 4), negatively affecting MS signal quality and sensitivity. Although the number of miscleavages decreased with statistical significance, quantitative depth did not improve as assessed by median peptide length distribution, Pearson correlation to bulk, dynamic range, and peptides per protein (Figure 3c, Suppl. Table 1, Suppl. Figure 1c,2c). Notably, the use of a reducing agent alone did not improve any assessing factors (Figure 3a-d, Suppl. Table 1). However, when extending the evaluation beyond simple identification counts, important biological relevance emerged. Despite a marginal decrease in total protein identifications, full reduction and alkylation enabled the unique detection of over 500 proteins involved in many cellular pathways, as revealed by gene ontology (GO) analysis (Figure 3e-g, Suppl. Figure 5). Additionally, subcellular localization analysis showed that reduction-only conditions introduced a bias toward proteins enriched in extracellular exosomes when compared to bulk samples (Figure 3g), suggesting altered sample representation. These findings highlight that while reduction and alkylation may introduce trade-offs in detection sensitivity and technical reproducibility, they also reveal biological information that would be missing otherwise. Thus, the inclusion of disulfide bond disruption steps should be considered carefully, based on the specific biological context and experimental goals.

**Figure 3.**
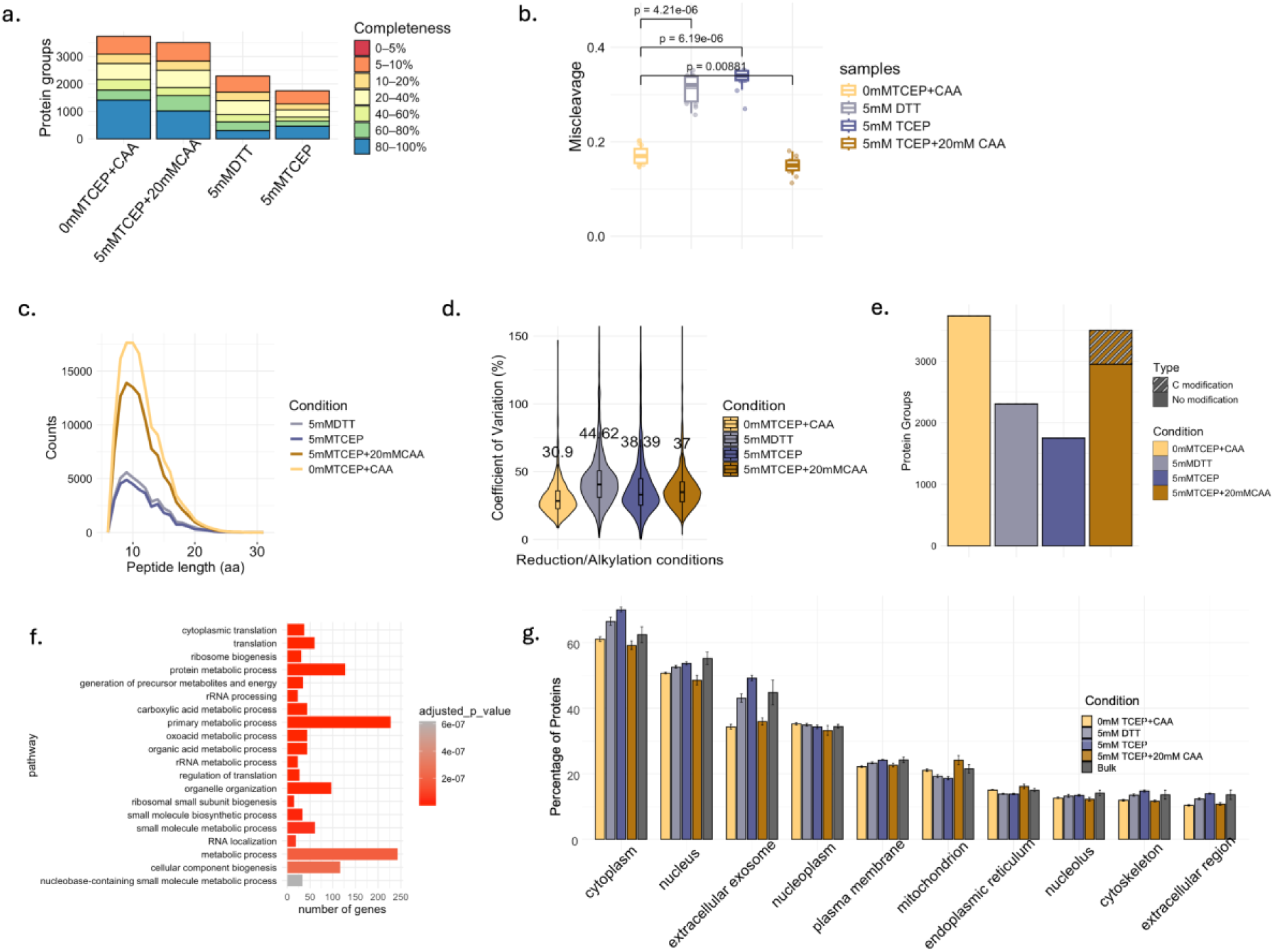
Effect of reduction and alkylation on single-cell proteome coverage and reproducibility. a. Protein coverage completeness across cells. Stacked bars show the number of protein groups binned by completeness (0–5% to 80–100%); b. Tryptic miscleavage of each digestion condition; c. Peptide length distributions (PSM versus amino-acid length) across different conditions; d. Coefficient of variation (CV) of protein intensities across cells shown with violin plots with median labeled for each condition; e. Total protein groups identified per condition (bars), with proteins detected only under reduction/alkylation highlighted (striped); f. Gene Ontology (GO) enrichment for pathways associated with proteins detected under reduction/alkylation; g. Subcellular localization of proteins identified from each condition.

### Simple evaporation control strategies for single-cell digestion

Due to the small reaction volumes necessary for single-cell proteomics workflows, evaporative loss presents a technical challenge, particularly in the absence of specialized dewpoint or humidity-controlled devices. Even under 100% humidity, buffer volumes in the low microliter range can experience substantial evaporation during incubation at 37°C. To address this, we increased the overall reaction volume by adding 0.5 µL of water prior to digestion for 4 hours under <=95% humidity using a standard 37°C tissue culture incubator (INC vs INC+0.5µl H_2_O) using A549 cells (n=15/condition). We observed an increase in protein coverage completeness, peptides per protein, bulk correlation, and peptide intensity distribution in comparison to digestion without the addition of water (Figure 4a, 4c, Suppl. Table 1). Despite equivalent concentrations of trypsin, the total volume loss with the addition of 0.5µL of water was 10% over 4 hours of trypsin digestion compared to 30% without the addition of water, indicating that a small increase in volume substantially reduces relative evaporative loss and stabilizes digestion.

**Figure 4.**
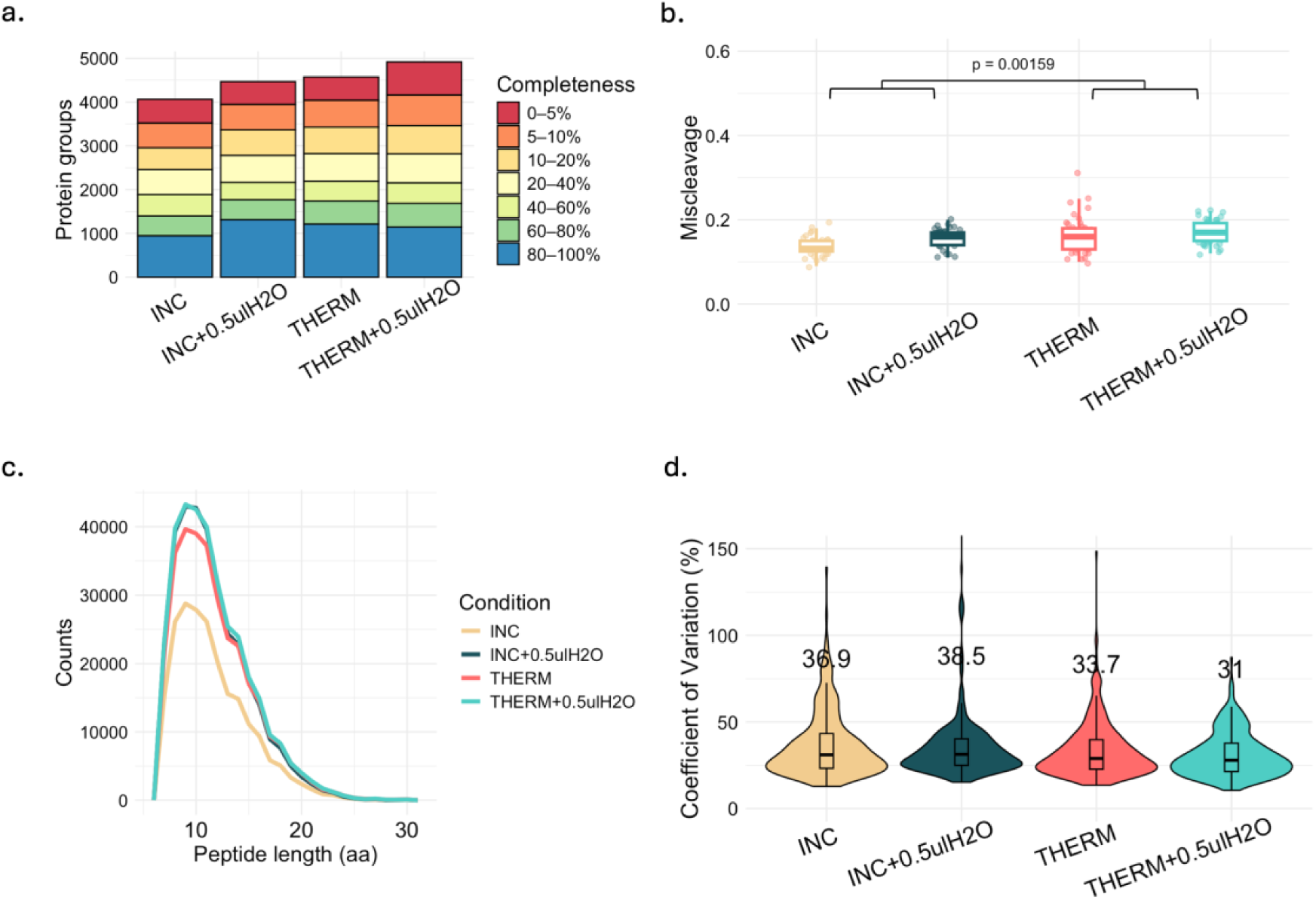
Effect of digestion conditions on single-cell proteome coverage and reproducibility. a. Protein coverage completeness across cells. Stacked bars show the number of protein groups binned by completeness (0–5% to 80–100%); b. Tryptic miscleavage of each condition; c. Peptide length distributions (PSM versus amino-acid length) across different conditions; d. Coefficient of variation (CV) of protein intensities across cells shown with violin plots with median labeled for each condition.

To determine whether comparable performance could be achieved without humidity control and provide an alternative option, we next evaluated a second method of evaporative control by performing digestion in a PCR thermal cycler with the lid heated to 42°C (THERM vs THERM+0.5µl H_2_O). We observed a similar volume loss. Miscleavage analysis indicated better performance in the cell culture incubator conditions (INC and INC+0.5µl H_2_O) compared to thermocycler incubation (THERM and TERM+0.5µl H_2_O) with p = 0.00159 (Figure 4b). Additionally, thermocycler-incubated samples exhibited significant lower CVs (p = 5.37 x 10^-5^), particularly when 0.5 µL water was added, suggesting improved reproducibility and data quality (Figure 4d). Despite these differences, the overall performance gain was comparable between incubator and thermocycler, enabling laboratories without a dedicated thermocycler to implement single-cell sample preparation using an incubator. More importantly, the addition of a small volume of water prior to digestion is an effective, accessible, and low-cost strategy to mitigate evaporation, thereby improving reproducibility and data consistency in single-cell digestion workflows without the need for dedicated thermocyclers.

### Four-Hour Trypsin Digestion as an Optimal Compromise for SCP

We next evaluated the effect of trypsin incubation time on protein digestion efficiency using 1ng of trypsin in 1ul of TEAB with 0.03% DDM with 0.5ul of water addition. A549 (n=15/condition) were used per condition. Reproducibility was higher between 4-16 hours of digestion, as measured by higher protein coverage completeness (∼1400 protein groups for 4-16h vs ∼1100 for 1-2h at 80-100% coverage) (Figure 5a, Suppl. Table 2). Longer digestion time also showed a decreasing trend in miscleavage, indicating better tryptic performance (Figure 5b). The distribution of peptide lengths, specifically those around the 7-13 amino acid length range, were time sensitive (Figure 5c). We observed fewer peptides at extended digestion times (i.e., 16h), perhaps as a result of over-digestion^20^. In contrast, shorter digestion times (i.e., 1h and 2h) showed evidence of incomplete proteolysis (Figure 5b, 5c), resulting in reduced peptide detection and higher CV values (Figure 5d). Overall, we identify that a digestion time of 4 hours yields high quality data, providing a balance between more completed trypsin digestion and minimize sample loss and negating the need for rapid-acting trypsin or specialized incubation equipment designed for single-cell workflows.

**Figure 5.**
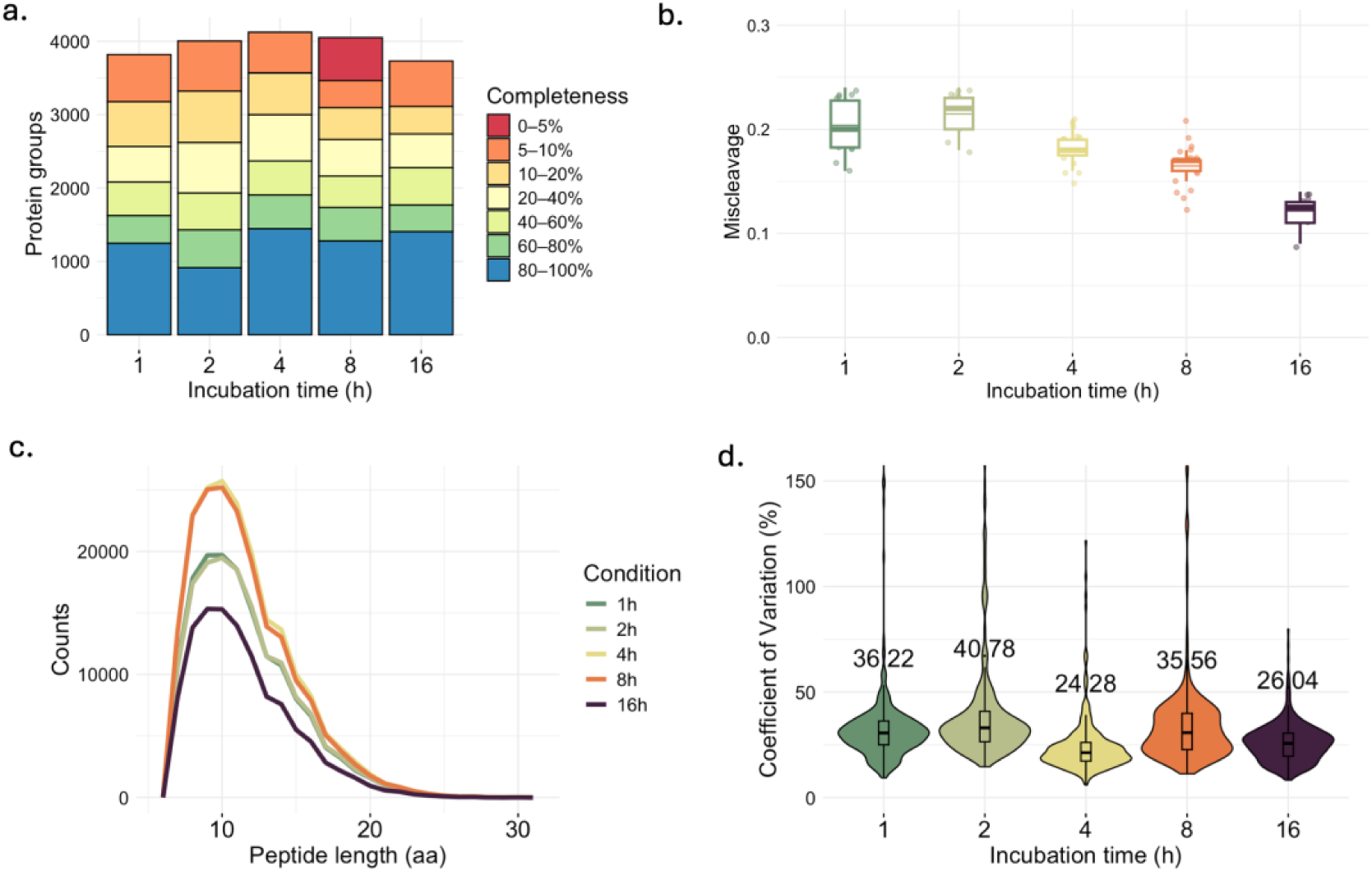
Effect of trypsin incubation time on single-cell proteome coverage and reproducibility. a. Protein coverage completeness across cells. Stacked bars show the number of protein groups binned by completeness (0–5% to 80–100%); b. Tryptic miscleavage of each incubation condition; c. Peptide length distributions (PSM versus amino-acid length) across different incubation times; d. Coefficient of variation (CV) of protein intensities across cells shown with violin plots with median labeled for each condition.

### Simple Plate and Digestion Choices Are Sufficient for Robust SCP

We also benchmarked plate format and trypsin proteolysis design. Comparing standard 96-well plates to low-bind 384-well plates under identical single-cell digestion conditions, 96-well plates yielded more than 300 proteins in the 80-100% completeness range, worse CVs (p = 2.08 x 10^-87^), and reduced peptide coverage, indicating that low-bind 384-well formats provide modest but consistent gains in depth and reproducibility (Suppl. Figure 6). In parallel, we evaluated double digestion (two additions of trypsin/LysC) and two-step versus one-step workflows and found no compelling benefit over simpler designs: double digestion did not improve peptide-to-protein ratio or bulk correlation, whereas single digestion yielded higher coverage completeness, fewer missed cleavages, lower CVs, improved peptide length distribution, and broader dynamic range (Suppl. Figure 7). Likewise, one-step and two-step protocols produced comparable overall quality with the one-step workflow yielding lower CVs and therefore more reproducible results (Suppl. Figure 8).

### Bioinformatics

Using our streamlined single-cell proteomics processing workflow, we next sought to evaluate the biological insights that may be gleaned from single-cell data. We analyzed four distinct cell types processed independently: immortalized A549, immortalized HEK-293, primary human astrocytes, and primary human monocytes. Uniform Manifold Approximation and Projection (UMAP)^21^ reveals clustering of cells according to tissue origin, consistent with tissue-specific proteomic signature (Figure 6a). Within each cluster, we were able to observe cell-selective marker expressions, including GFAP (astrocytes), CAC1H marker (hCaV3.2), CAC1G marker (hCaV3.1), ARG2 marker (A549) and CD14 marker (monocytes) (Figure 6b)^22^. Remarkably, we were also able to detect intra-cluster differences within parental HEK-293 cells, and HEK-293 cells transfected with Cav3.1T-type or Cav3.2 T-type calcium channels (Figure. 6c). To further evaluate the robustness, we performed analysis using PC12 cells from mice, UMAP distinguished undifferentiated (day 0) from differentiated states and suggested a graded trajectory from day 2 to day 6 (Figure 6d).

**Figure 6:**
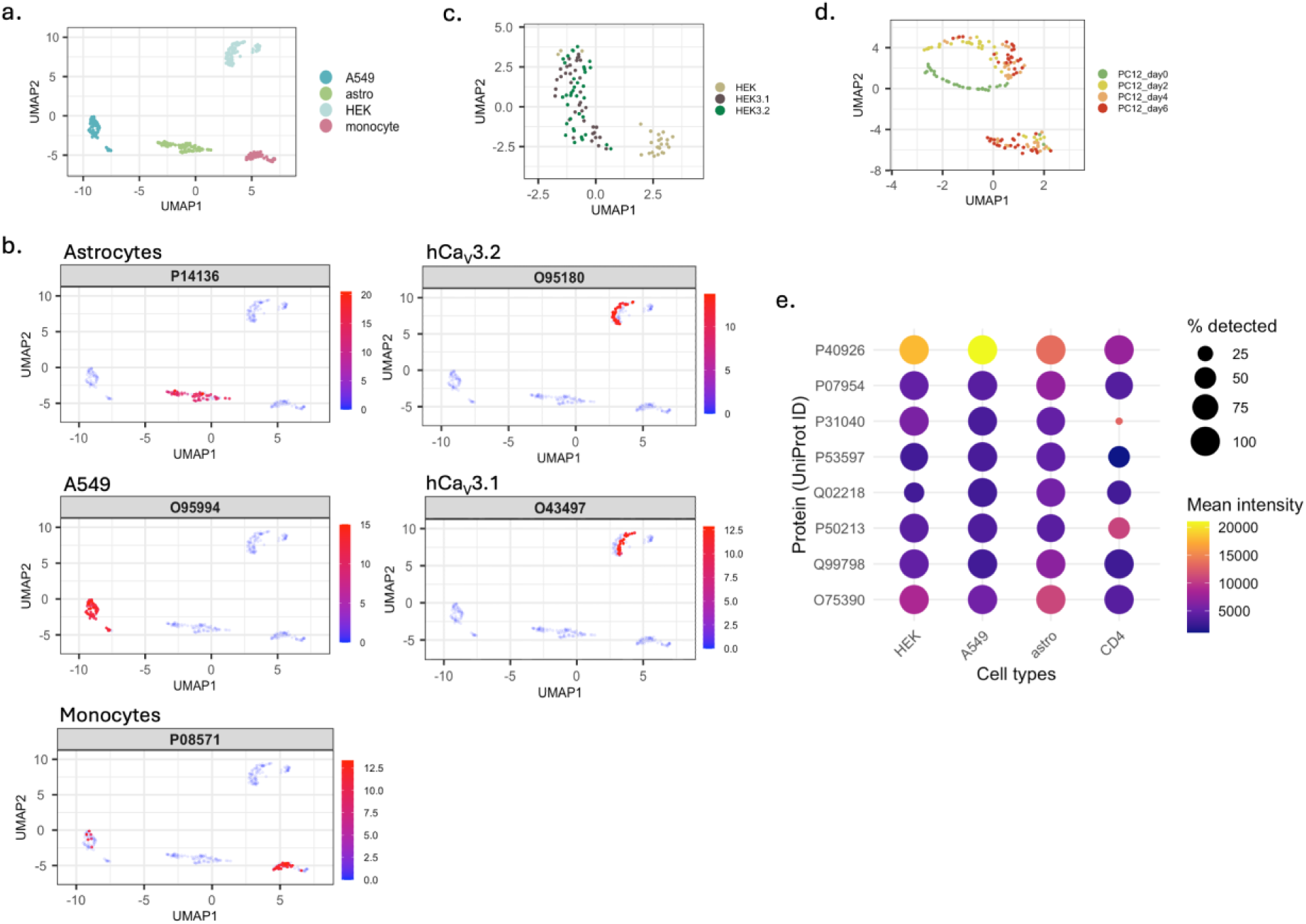
Bioinformatics analysis of the single-cell sample acquired with the miniaturized sample preparation method. a. Clustering of A549 (turquoise), astrocyte (green), HEK (light blue), and T cell (pink) on UMAP; b. Protein markers, GFAP (P14136) for astrocytes, CAC1H (O95180) for hCaV3.2, CAC1G (O43497) for hCaV3.1, ARG2 (O95994) for A549 and CD14 (P08571) for monocytes, highlighted on UMAPs; c. UMAP clustering untransfected HEK-293F (light brown), transfected hCa_V_3.1 (dark brown), and transfected hCa_V_3.2 (green); d. UMAP clustering undifferentiated PC12 (green), day 2 differentiated PC12 (yellow), day 4 differentiated PC12 (orange) and day 6 differentiated PC12 (red); e. The protein coverage completeness of the TCA cycle in HEK, A549, astrocyte and monocytes presented on dot plot.

To assess reproducibility across various cellular contexts, we evaluated the protein coverage completeness for components of the Tricarboxylic Acid cycle, a metabolic pathway common to the cellular contexts in this study (Figure 6e). We observed a majority of proteins with 100% coverage, with the exception of succinate dehydrogenase (subunit A) (Uniprot ID: P31040). This might reflect low abundance, or binding to cofactors that occlude digestion without stringent denaturation^23^.

Together, these findings suggest that our streamlined single-cell proteomics workflow delivers high-quality and reproducible data from which biologically meaningful insights may be derived from heterogeneous populations.

## Discussion

Here we provide a systematic evaluation of single-cell proteomics (SCP) processing parameters and demonstrate that high-quality proteomic data may be generated without the use of highly specialized laboratory equipment, thus broadening access to SCP technologies. Our robust, label-free sample preparation protocol requires only a single-cell dispenser, conventional incubation chamber, and easily accessible well plates.

In this streamlined workflow, which uses only commercial 384-well plates and a regular incubator, we were able to define a small group of parameters that primarily govern data quality. For enzyme load, increasing trypsin beyond 1ng for standard cells of 10-15µm sized reduced miscleavage but did not improve, and sometimes degrade, quantitative depth by shifting signals towards trypsin-derived peptides and reducing detection of analytically useful peptides. Conversely, lowering the trypsin amount led to incomplete proteolysis and noisier quantification. Technical choices around evaporation control and plate format had more modest but consistent effects. Simple intervention of water supplementation and either a humidified chamber or a thermocycler with heated lid was sufficient to reduce volume loss and stabilize quantitative metrices without requiring specialized dewpoint-controlled devices. The value of reduction and alkylation appeared to be context dependent. On the timsTOF platform, omitting these steps improved reproducibility and reduced reagent-derived contamination for routine measurements, yet full reduction/alkylation uniquely revealed subsets of disulfide-rich and metabolically relevant proteins.

While tuning enzyme load, digestion conditions, and evaporation control improve performance, we further highlight that an equally important question is how SCP workflows are evaluated. When the field of bulk proteomics was developing, the number of proteins one could identify was the metric that everyone chased - the more proteins one could identify, the better^24^. Indeed, this still applies to the development of new instrumentation^25,26^ and new methodology^27,28^. The same has been true with single-cell proteomics to date. However, the number of proteins identifiable is far less important than the number of proteins that can be reproducibly quantified across all of the cells being analyzed. Building on this perspective, we propose a comprehensive and more biologically meaningful approach to evaluate single-cell proteomics data by incorporating quantitative consistency and biological interpretability. We suggest four key dimensions of data quality assessment beyond protein counts: reproducibility, reliability, and quantitative accuracy. While these metrics may yield smaller numbers (e.g., the number of proteins that one can achieve 80% completeness is always going to be lower than the total identified), they provide a more realistic view of the proteome that can be robustly inferred. If it is important to focus on a single metric to simplify a message, then protein coverage completeness or peptide metrics are the most meaningful.

In line with this framework, we advocate against over-reliance on bulk-style differential expression tools, such as volcano plots and group-level enrichment analyses, when analyzing SCP data. These approaches assume population-level homogeneity and risk obscuring the very heterogeneity that single-cell methods are designed to reveal. Instead, we suggest focusing on cell-resolved visualization strategies such as probabilistic principal component analysis (PPCA), Uniform Manifold Approximation and Projection (UMAP), heatmaps, and individual protein-level tracking to uncover meaningful variability and subpopulation structure.

Looking ahead, the accessible and adaptable SCP workflow presented here provides a solid foundation for integration with other single-cell modalities. Coupling proteomic data with single-cell transcriptomics, lipidomics, metabolomics, spatial profiling, or patch-seq could yield multidimensional insights into cellular function and disease mechanisms. As mass spectrometry sensitivity and computational analysis tools continue to advance, prioritizing quantitative fidelity, data reproducibility, and biological interpretability will be essential for transforming single-cell proteomics from a technical frontier into a translationally impactful field.

## Acknowledgments

We thank BioRender for support in creating the figure illustrations. We also thank the Proteomics Core Facility at Life Science Institution, University of British Columbia for assisting in data acquisition. A.E. thanks Oregon State University ‘s Department of Chemistry for a Summer Internship Fellowship. This work was supported in part by an OSU/HP collaboration grant to C.M. MAM is the Terry Fox Leader in Cancer Genome Science.

## Data and code availability

All data and code are available in public repository. The mass spectrometry raw data have been deposited to ProteomeXchange via MassIVE (MSV000100666). Associated code for the data analysis is included in R markdown or R script files at GitHub repository: https://gist.github.com/lucychii/8af478f7e4a7075372338264a7d5af1a.

## Supplementary information

**Supplementary Figure 1.**
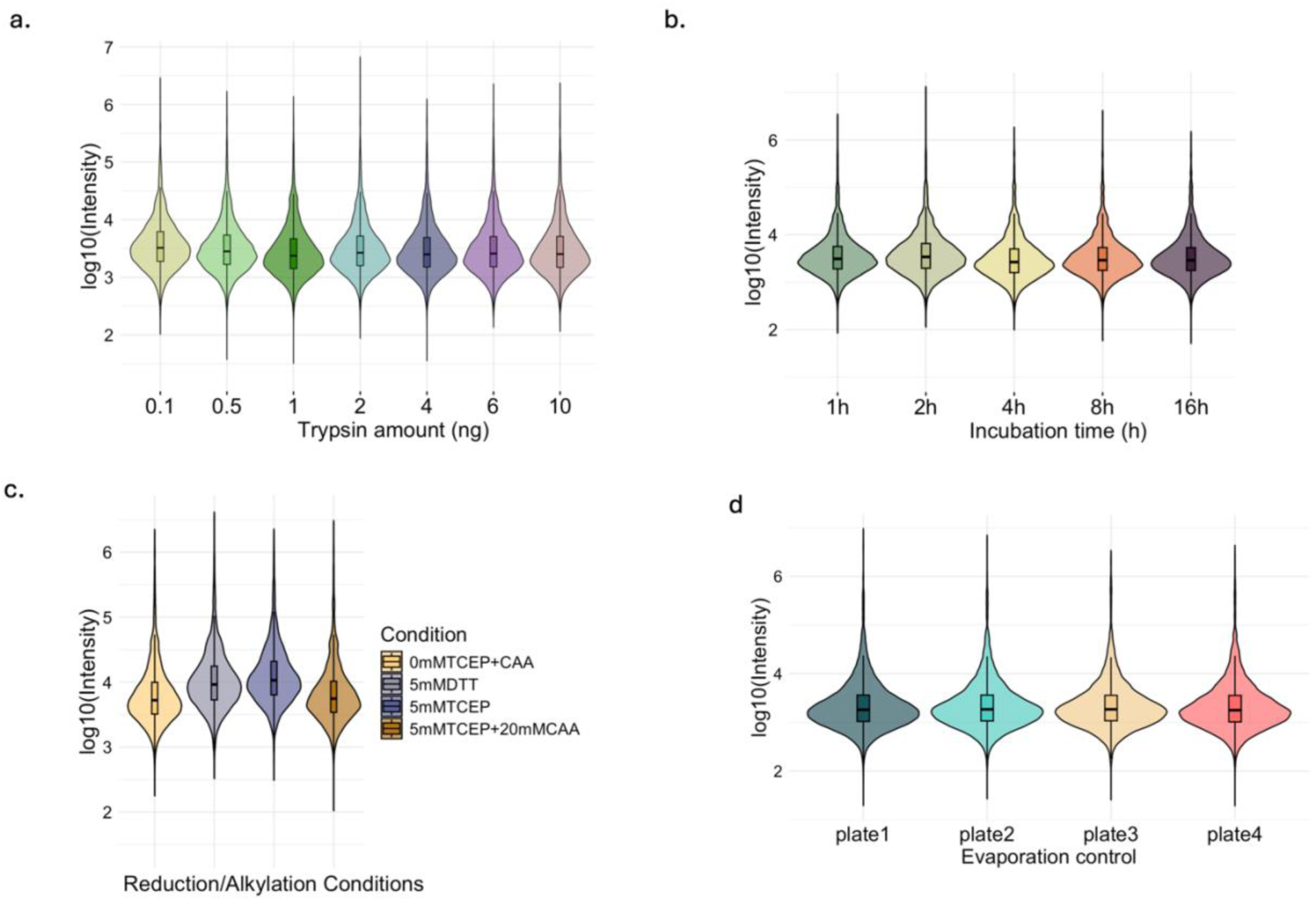
Dynamic range of protein intensity across sample preparation parameters. Log_10_ transformed protein intensity distributions presenting dynamic range across different sample preparation conditions. Each distribution indicates the intensity spread and the central tendency, enabling comparison of overall protein trend and dynamic range across workflows.

**Supplementary Figure 2.**
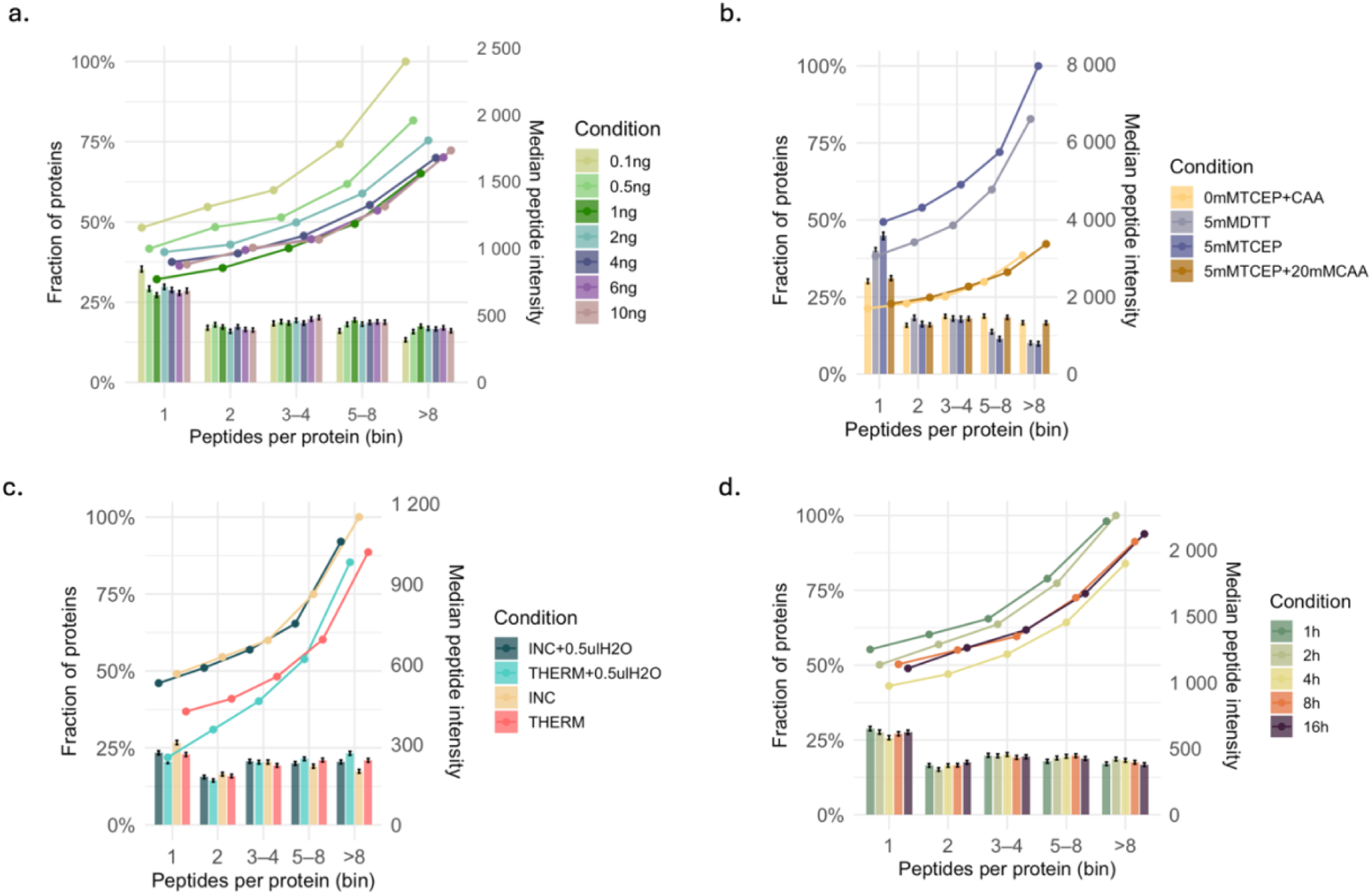
Peptide-per-protein ratio depth and median peptide intensity by condition. Bar plots show the average number of peptides identified per protein across conditions and the overlaid dots indicate the corresponding median peptide intensity.

**Supplementary Figure 3.**
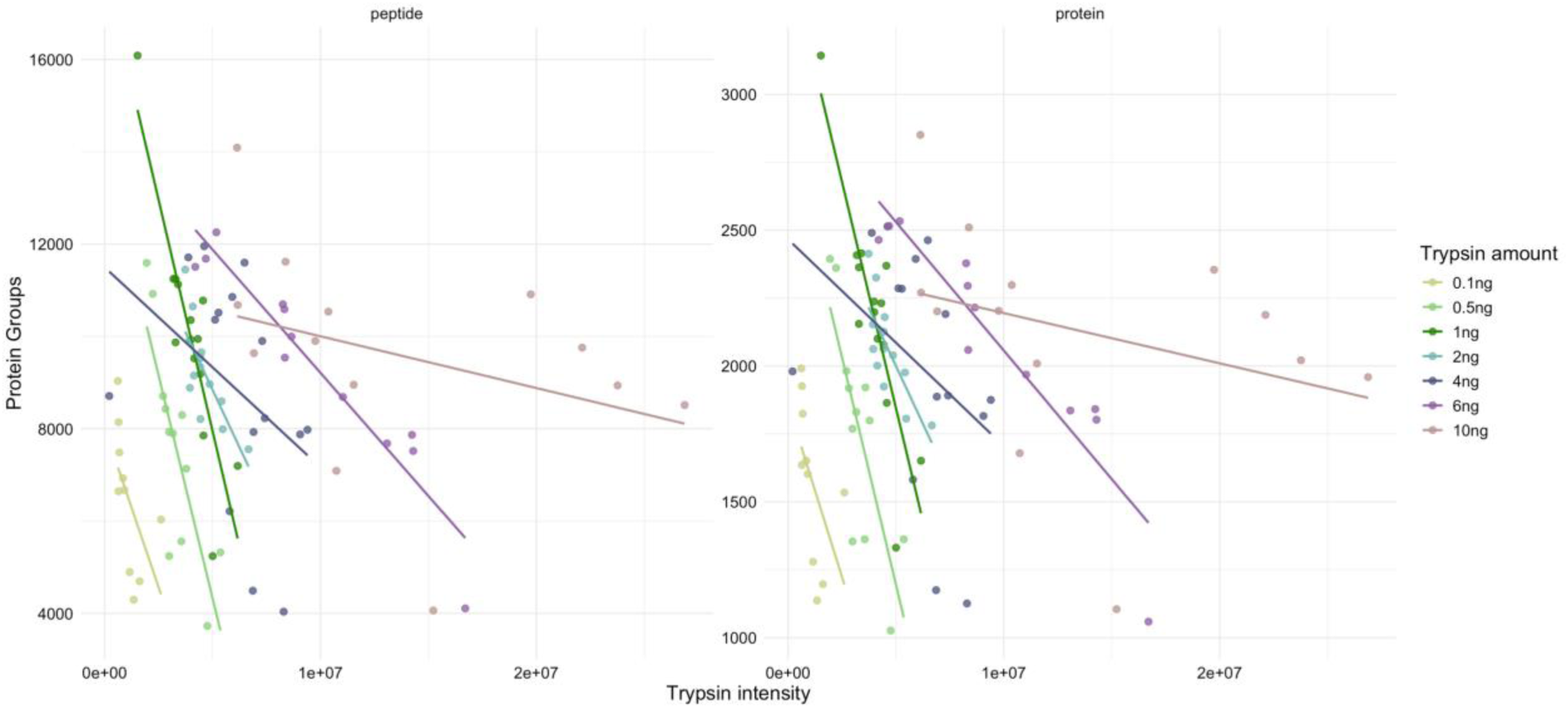
Relationship between trypsin intensity and the number of protein IDs (right) and peptide IDs (left) for each condition.

**Supplementary Figure 4.**
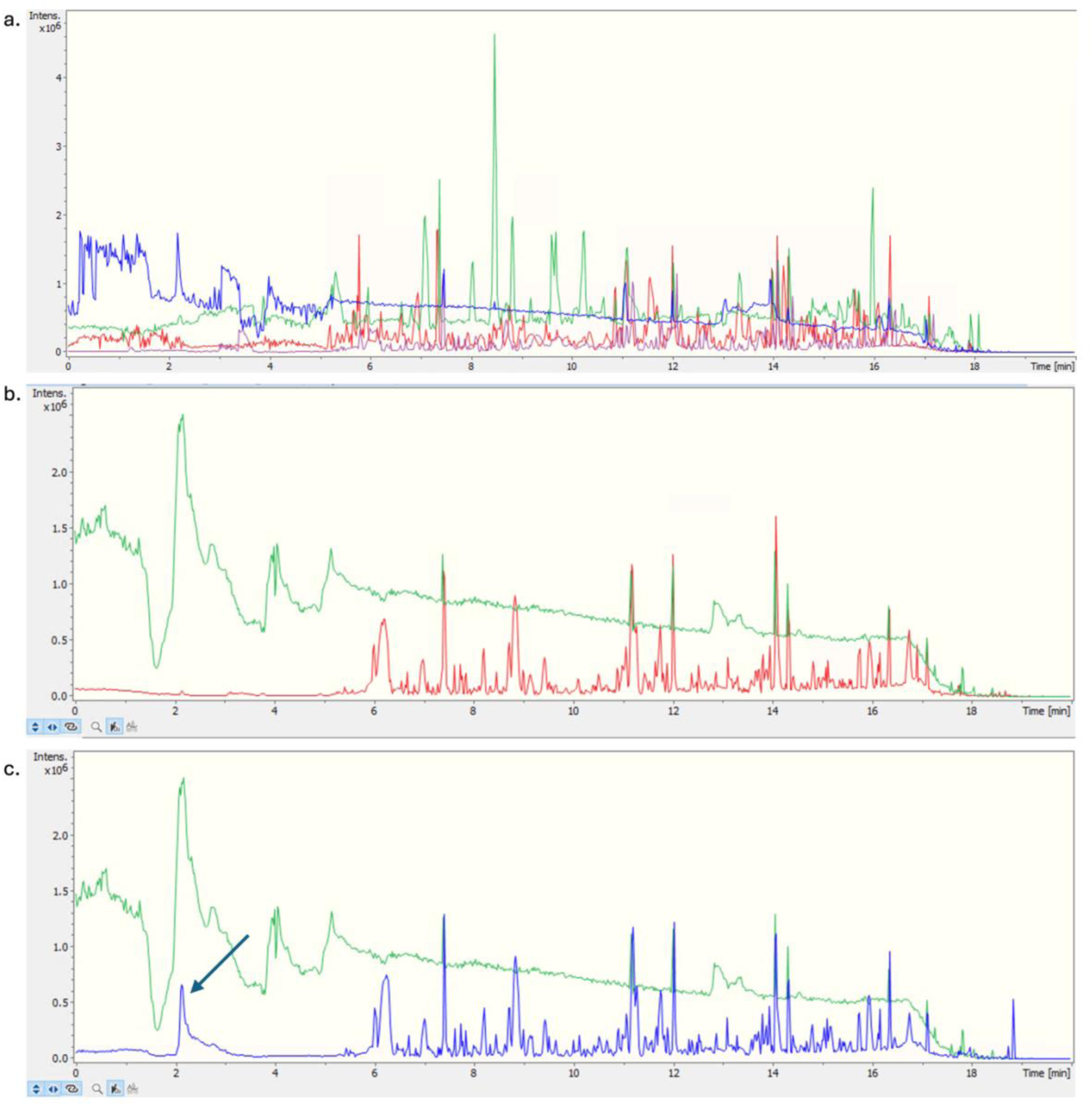
Chromatograms for reduction and alkylation condition tests. a. An overlap view of the chromatogram for 0mMTCEP+CAA (purple), 5mM TCEP+20mM CAA (blue), 5mM TCEP (green) and 5mM DTT (red); b-c. Chromatograms showing contamination carryover. Samples with reduction and alkylation of 5mM TCEP+20mM CAA shown in green; sample injection before running reduced and alkylated samples (red, panel b) compared to one after running reduced and alkylated samples (blue, panel c).

**Supplementary Figure 5.**
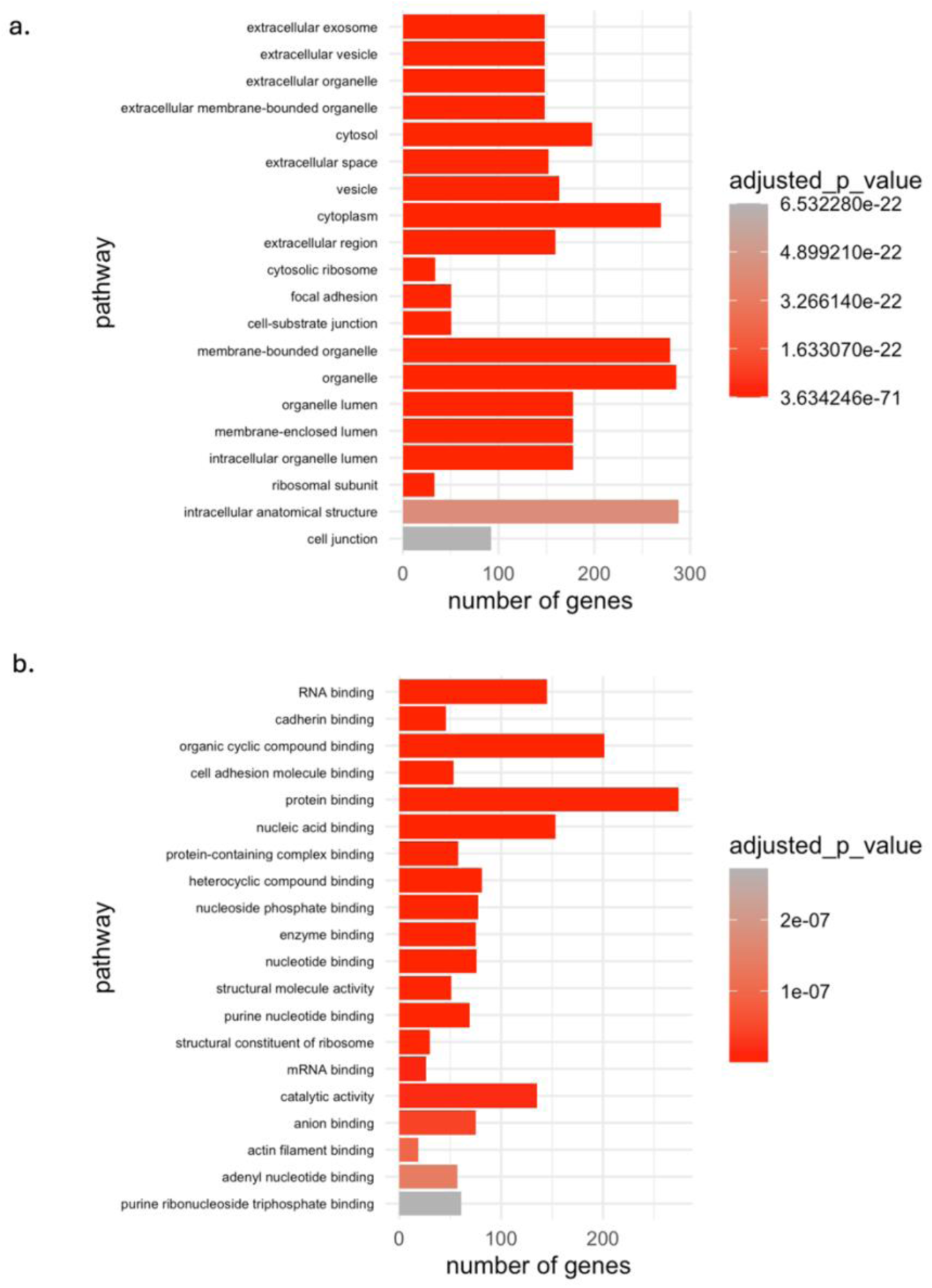
GO analysis of proteins identified from reduction and alkylation. a. Cellular component GO analysis; b. Molecular function GO analysis.

**Supplementary Figure 6.**
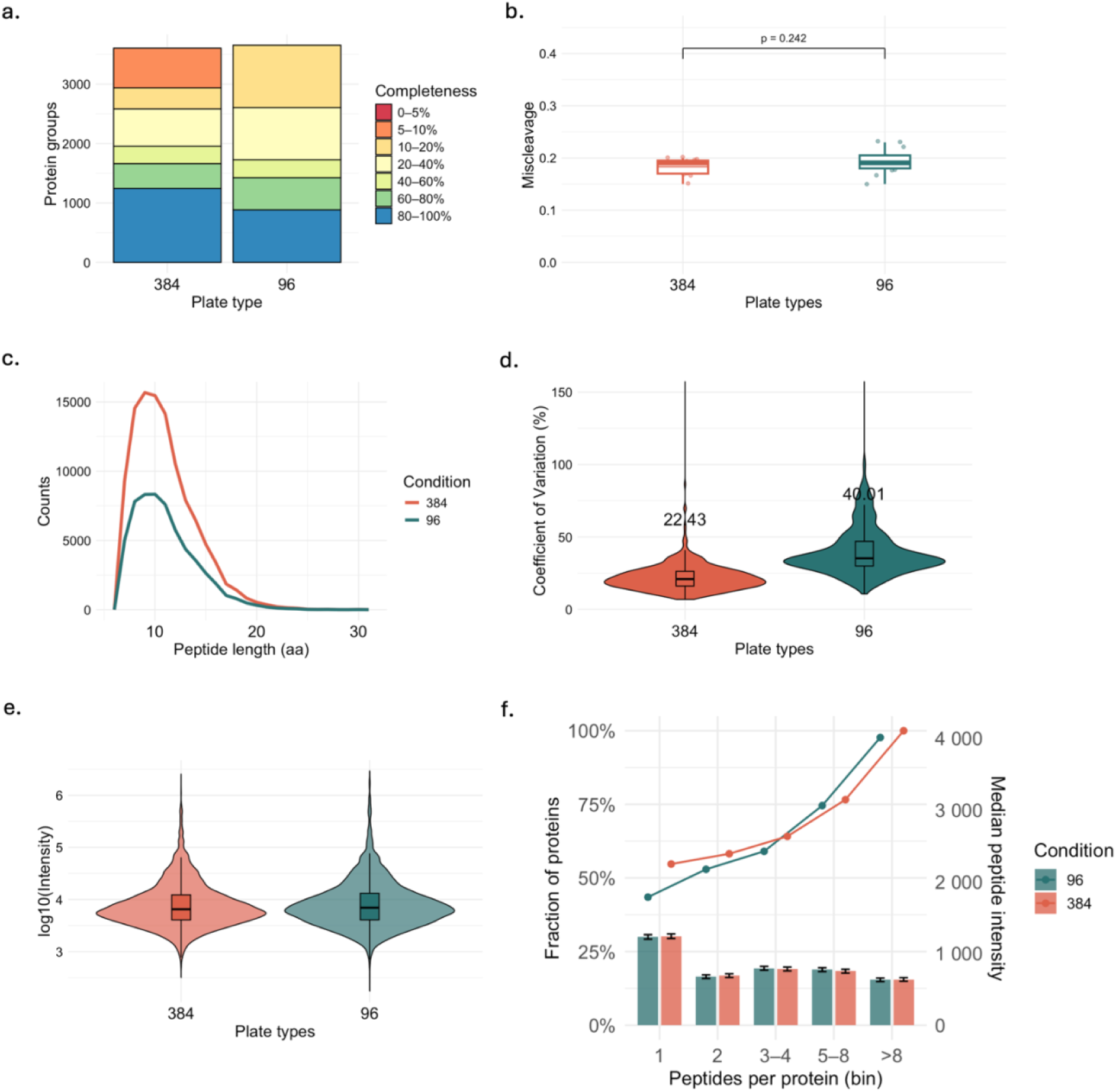
Effect of plate type (384 versus 96-well plate) on single-cell proteome coverage and reproducibility. a. Protein coverage completeness across cells. Stacked bars show the number of protein groups binned by completeness (0–5% to 80–100%); b. Tryptic missed-cleavage of each plate type; c. Peptide length distributions (PSM versus amino-acid length) across different plate types; d. Coefficient of variation (CV) of protein intensities across cells shown with violin plots with median labeled for each condition; e. Dynamic range of protein intensity between conditions; f. Peptide-per-protein ratio depth and median peptide intensity by conditions.

**Supplementary Figure 7.**
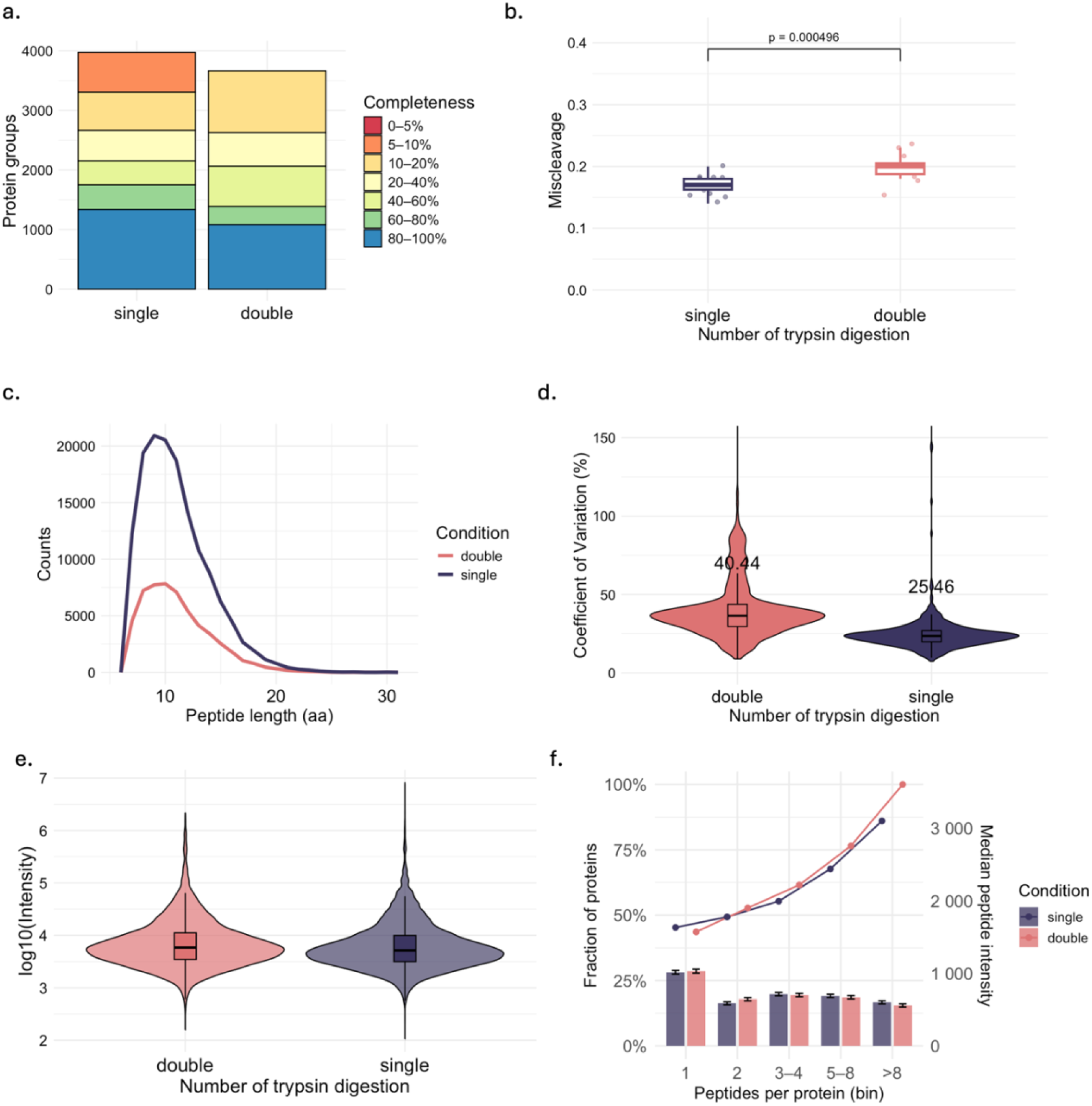
Effect of single versus double trypsin digestion on single-cell proteomics data quality. a. Protein coverage completeness across cells. Stacked bars show the number of protein groups binned by completeness (0–5% to 80–100%); b. Tryptic missed-cleavage comparing single versus double trypsin digestion; c. Peptide length distributions (PSM versus amino-acid length) across workflows; d. Coefficient of variation (CV) of protein intensities across cells shown with violin plots with median labeled for each condition; e. Dynamic range of protein intensity between conditions; f. Peptide-per-protein ratio depth and median peptide intensity by conditions.

**Supplementary Figure 8.**
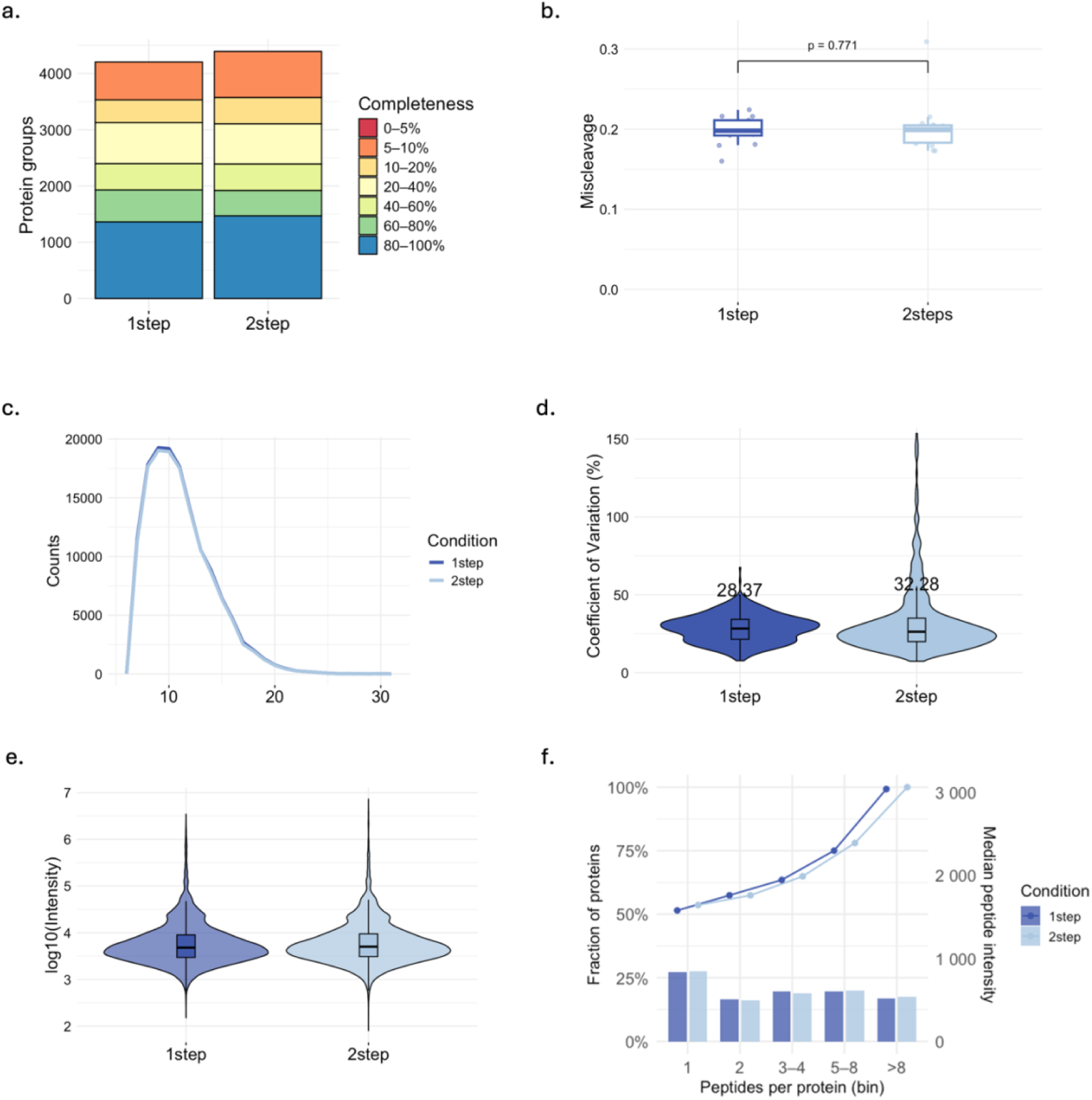
Effect of number of steps involved on single-cell proteomics data quality. a. Protein coverage completeness across cells. Stacked bars show the number of protein groups binned by completeness (0–5% to 80–100%); b. Tryptic missed-cleavage comparing 1-step versus 2-step protocols; c. Peptide length distributions (PSM versus amino-acid length) across workflows; d. Coefficient of variation (CV) of protein intensities across cells shown with violin plots with median labeled for each condition; e. Dynamic range of protein intensity between conditions; f. Peptide-per-protein ratio depth and median peptide intensity by conditions.

**Supplementary Figure 9.**
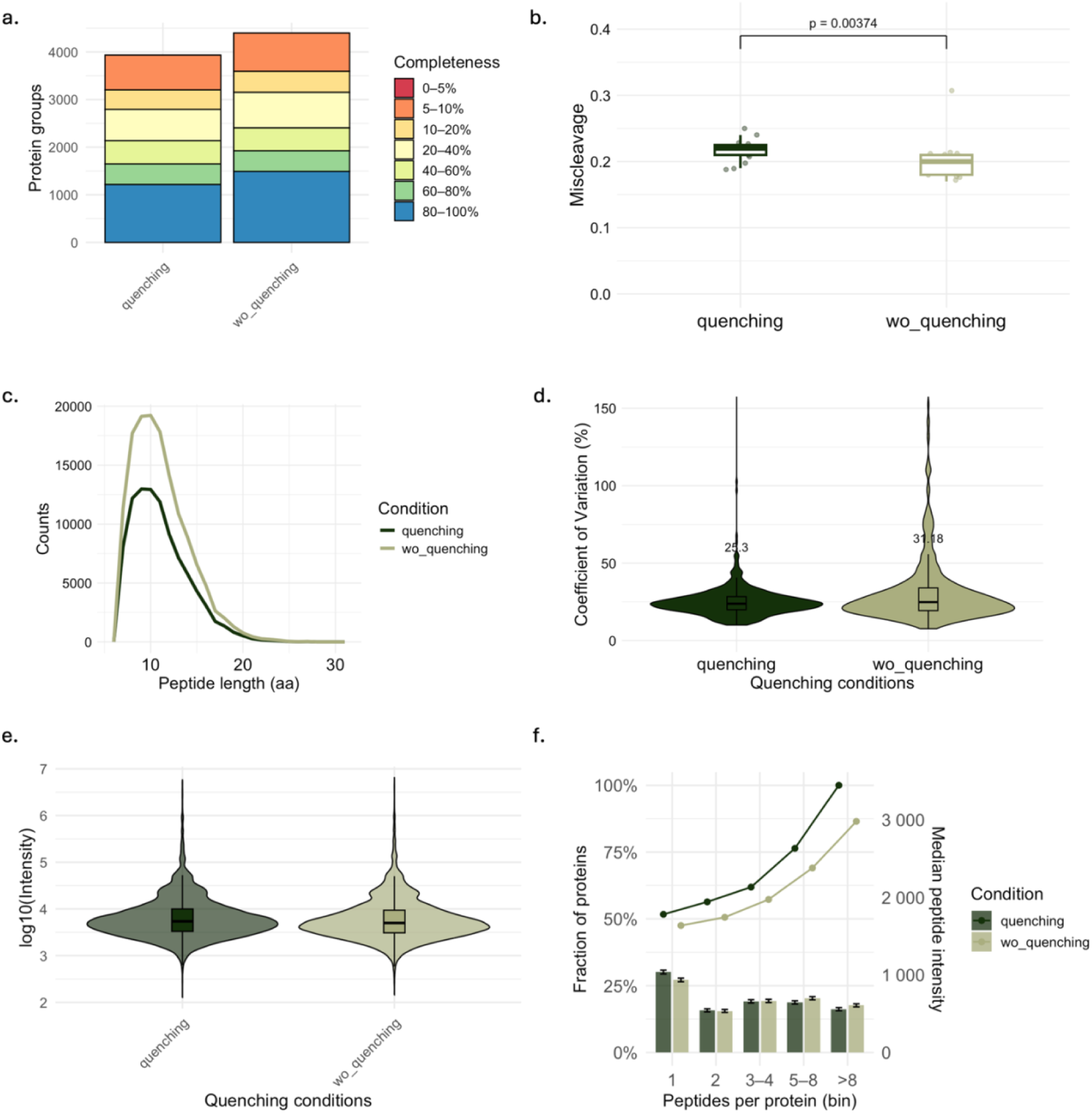
Effect of quenching on single-cell proteomics data quality. a. Protein coverage completeness across cells. Stacked bars show the number of protein groups binned by completeness (0–5% to 80–100%); b. Tryptic missed-cleavage comparing with and without quenching; c. Peptide length distributions (PSM versus amino-acid length) across workflows; d. Coefficient of variation (CV) of protein intensities across cells shown with violin plots with median labeled for both conditions; e. Dynamic range of protein intensity between conditions; f. Peptide-per-protein ratio depth and median peptide intensity by conditions.

**Supplementary Figure 10.**
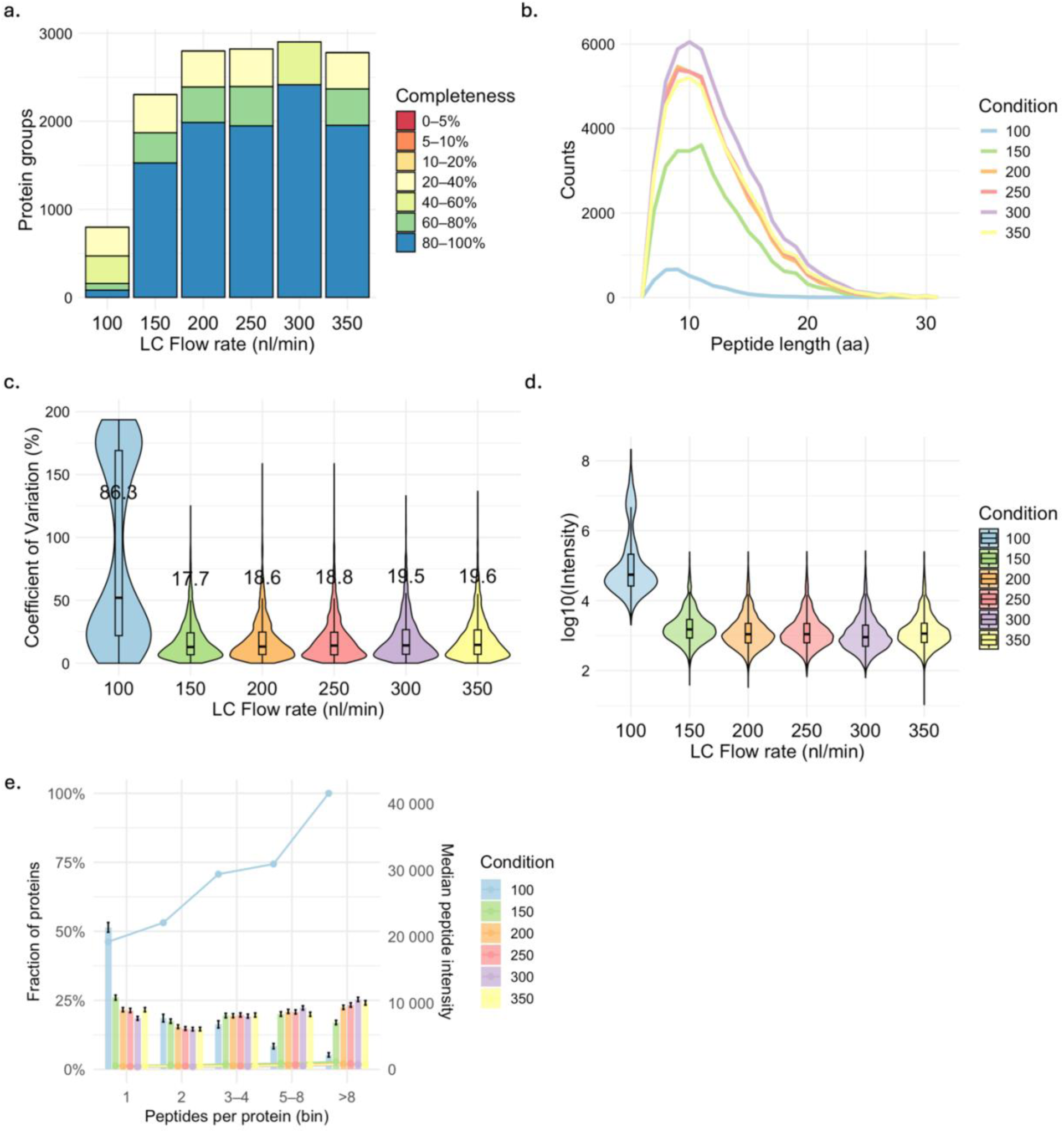
Effect of LC flow rate (15min gradient) coupled with dia-PASEF window MS with 1.28s cycle time (250pg Hela per injection). a. Protein coverage completeness across cells. Stacked bars show the number of protein groups binned by completeness (0–5% to 80–100%); b. Peptide length distributions (PSM versus amino-acid length) across workflows; c. Coefficient of variation (CV) of protein intensities across injections shown with violin plots with median labeled for both conditions; d. Dynamic range of protein intensity between conditions; e. Peptide-per-protein ratio depth and median peptide intensity by conditions.

**Supplementary Figure 11.**
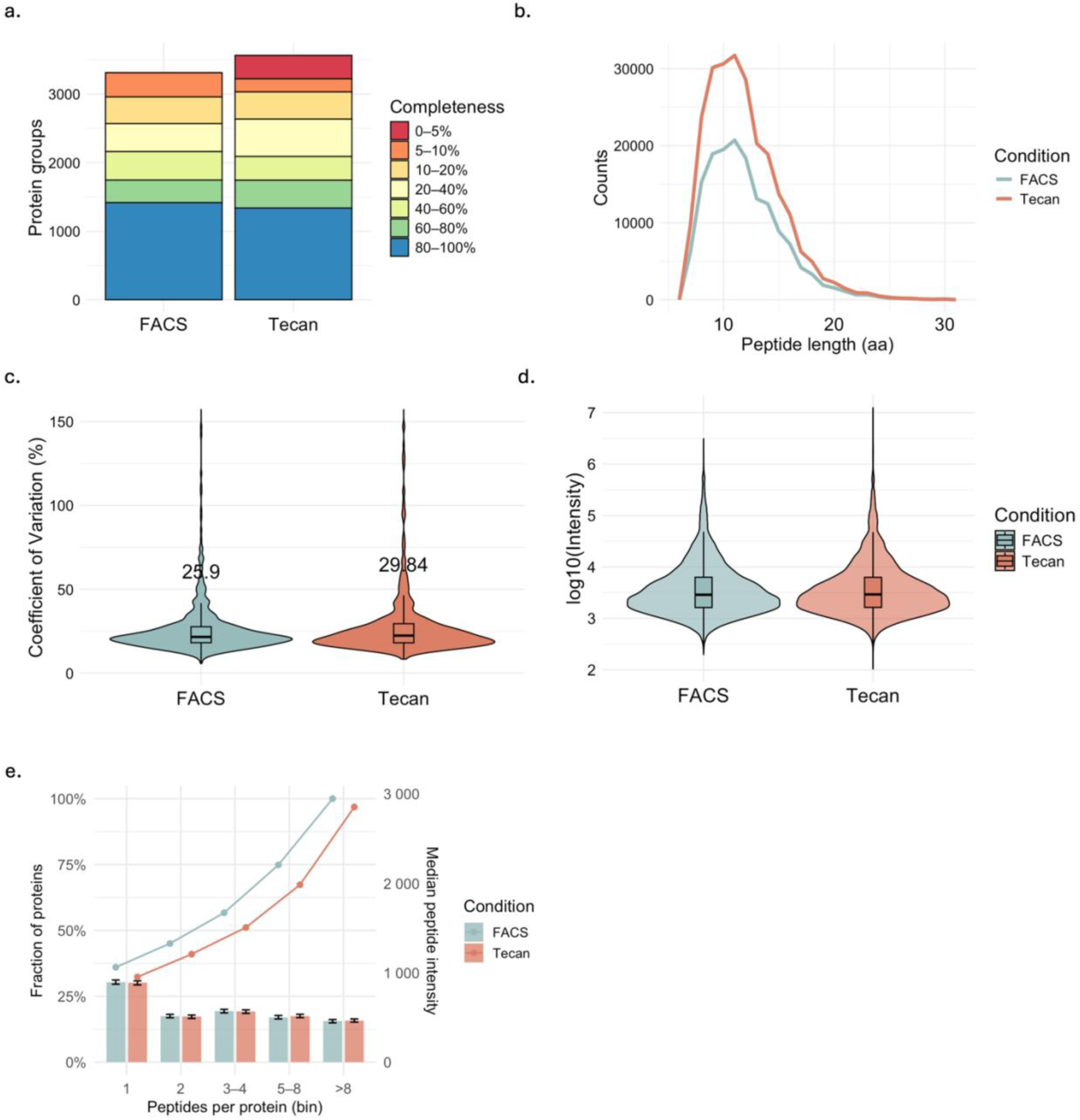
Difference between single cell dispensers: FACS vs Tecan Uno Dispenser. a. Protein coverage completeness across cells. Stacked bars show the number of protein groups binned by completeness (0–5% to 80–100%); b. Peptide length distributions (PSM versus amino-acid length) across workflows; c. Coefficient of variation (CV) of protein intensities across cells shown with violin plots with median labeled for both conditions; d. Dynamic range of protein intensity between conditions; e. Peptide-per-protein ratio depth and median peptide intensity by conditions.

**Supplementary Figure 12.**
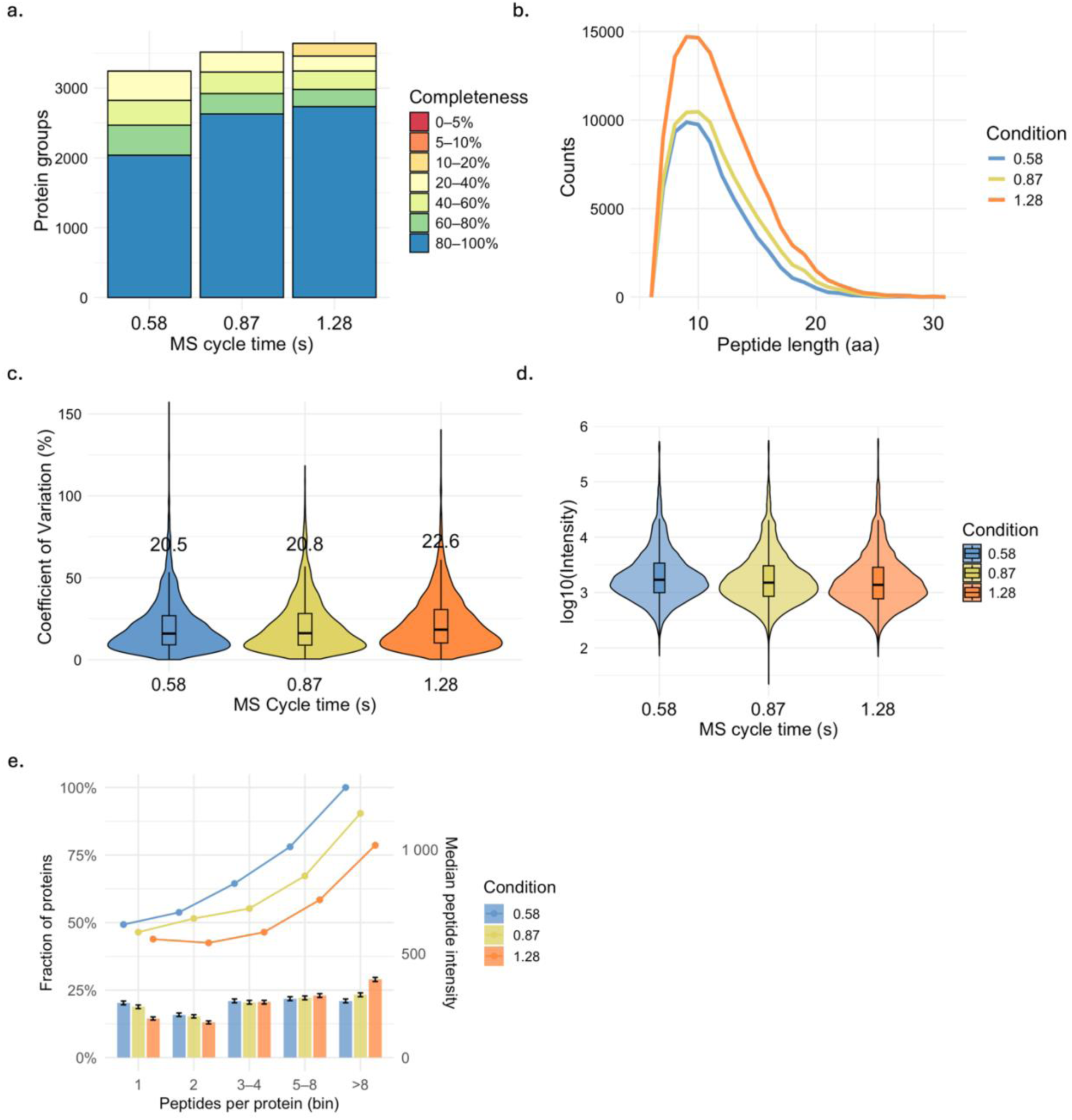
Effect of MS cycle time with 300nl/min flow rate and 15min gradient LC (250pg Hela per injection). a. Protein coverage completeness across cells. Stacked bars show the number of protein groups binned by completeness (0–5% to 80–100%); b. Peptide length distributions (PSM versus amino-acid length) across workflows; c. Coefficient of variation (CV) of protein intensities across injections shown with violin plots with median labeled for both conditions; d. Dynamic range of protein intensity between conditions; e. Peptide-per-protein ratio depth and median peptide intensity by conditions.

**Supplementary Figure 13.**
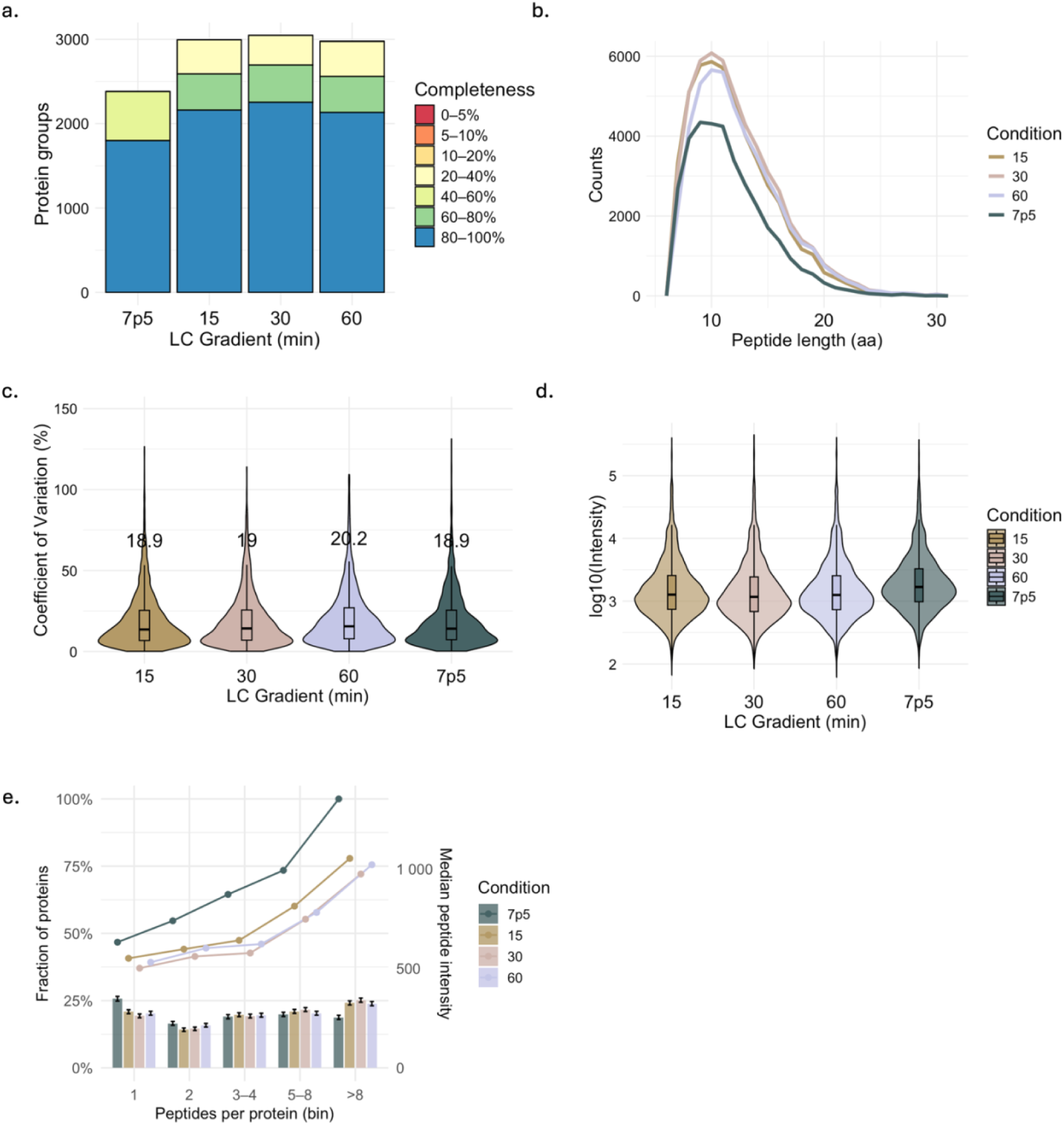
Effect of LC gradient (300nl/min flow rate) coupled with dia-PASEF window MS with 1.28s cycle time (250pg Hela per injection). a. Protein coverage completeness across cells. Stacked bars show the number of protein groups binned by completeness (0–5% to 80–100%); b. Peptide length distributions (PSM versus amino-acid length) across workflows; c. Coefficient of variation (CV) of protein intensities across injections shown with violin plots with median labeled for both conditions; d. Dynamic range of protein intensity between conditions; e. Peptide-per-protein ratio depth and median peptide intensity by conditions.

**Supplementary Figure 14.**
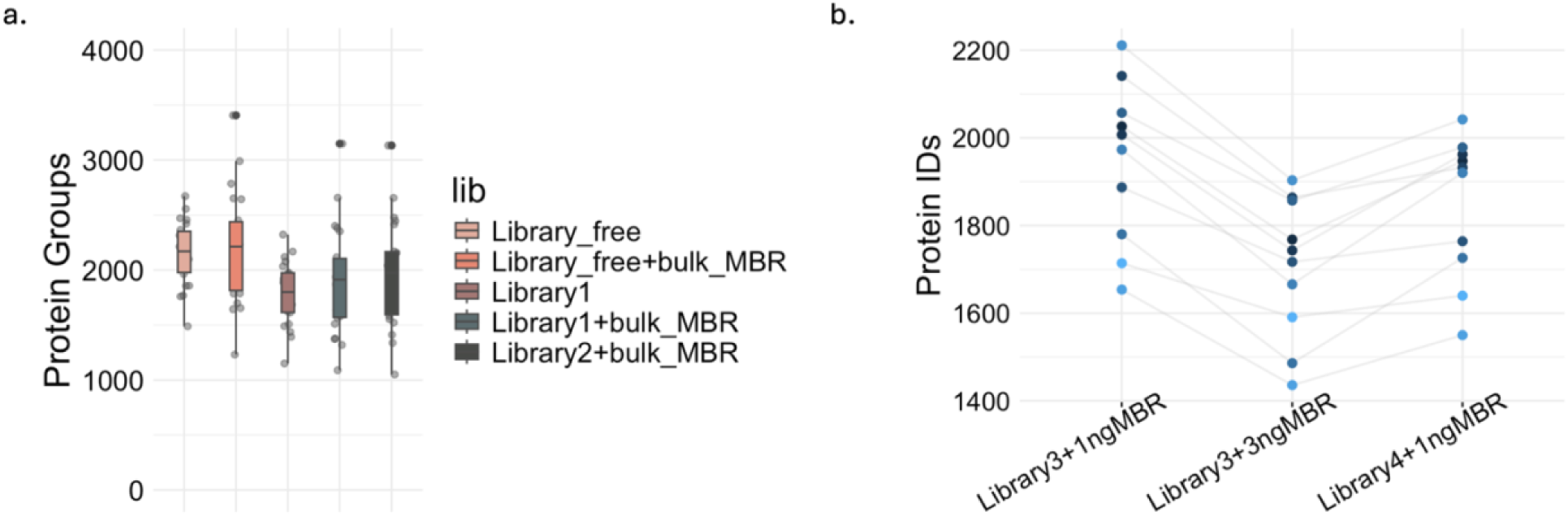
Effect of DIA-NN search strategies. a, Average number of protein groups identified from single A549 cells acquired using a 15 min LC gradient (300 nL/min) with dia-PASEF (1.28 s cycle time), comparing different library settings with and without bulk Match-Between-Runs. Library 1: larger library generated using a 30 min LC method. Library 2: smaller, method-matched library generated using the same LC-MS/MS conditions as the single-cell runs. b, Comparison of three DIA-NN search strategies for 250pg HeLa injections. Library 3: larger library generated on timsTOF Ultra 2. Library 4: smaller, method-matched library generated using the same LC–MS/MS conditions as the 250pg HeLa runs.

**Supplementary Table 1.**
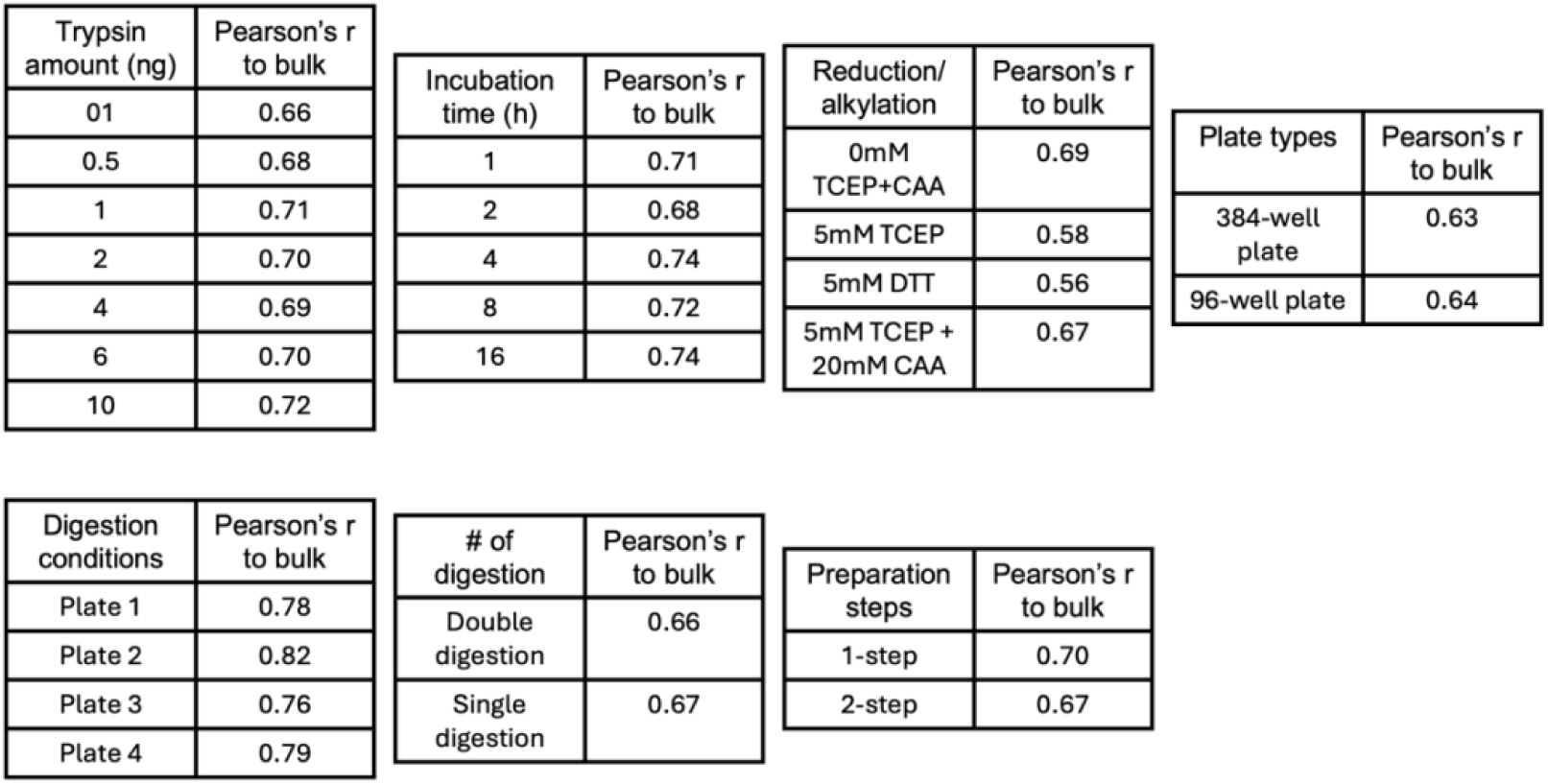
Pearson correlation of protein intensity between bulk and single cell samples for each condition tested. For each comparison, the mean intensity across all single cell samples is calculated and compared to that of the bulk samples.

**Supplementary Table 2.**
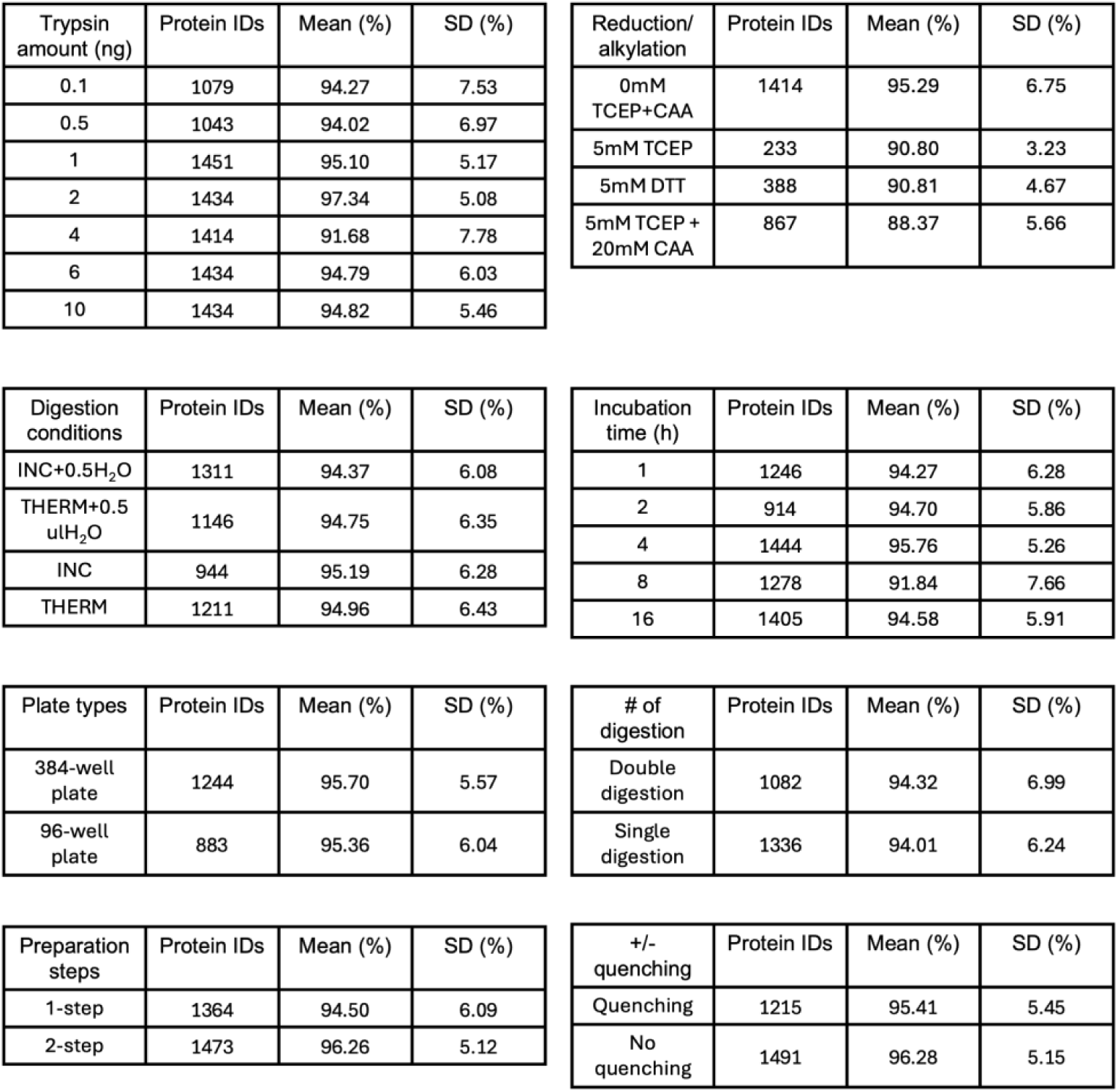
The protein counts, mean and standard deviation for protein coverage completeness of 80-100% for each condition.

**Supplementary Table 3.**
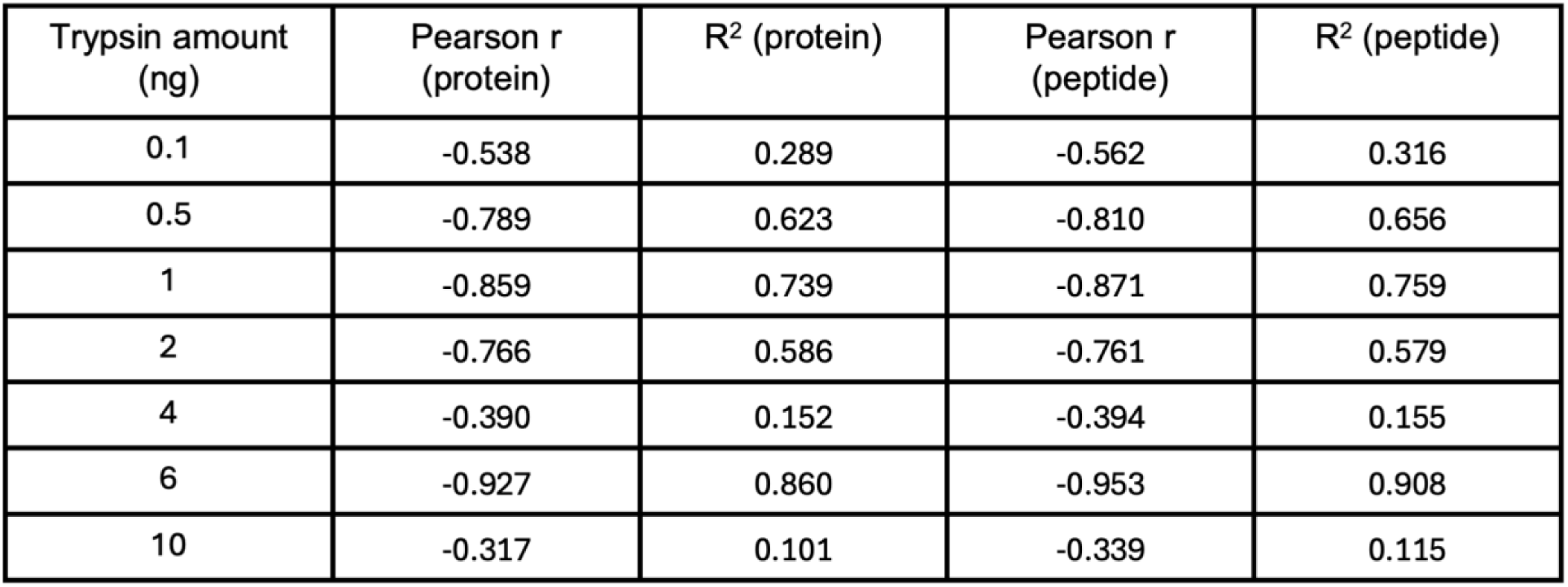
Pearson correlation r values and R^2^ values for each condition plotted in Suppl. Figure 3.

