## Supplementary Table 2 for "Bridging Simplicity and Depth in Single-Cell Proteomics: A Cost-Effective Workflow and Expanded Framework for Data Evaluation"

Supplementary Table 2. The protein counts, mean and standard deviation for protein coverage completeness of 80-100% for each condition.

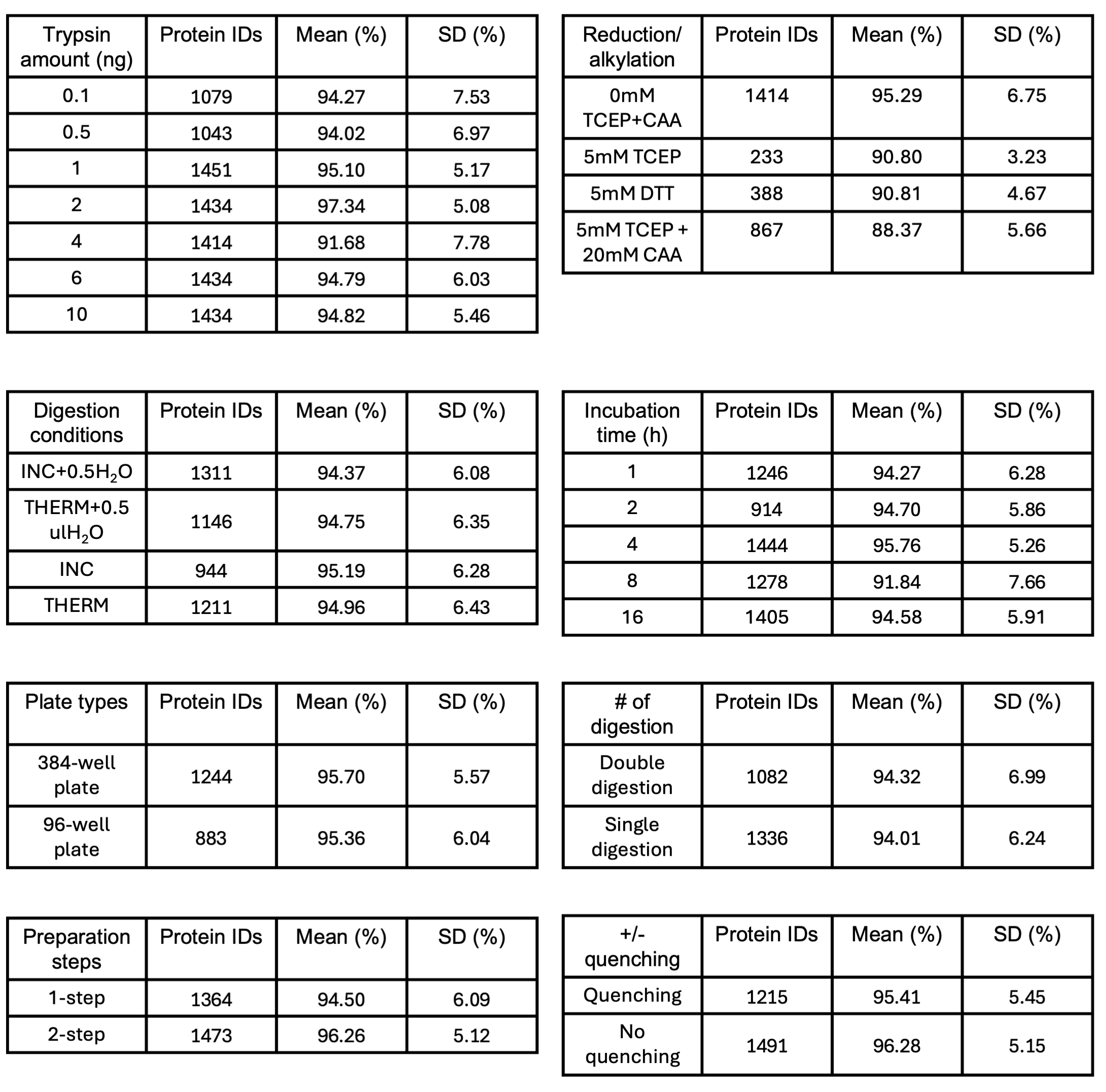
