## Supplementary Table 1 for "Bridging Simplicity and Depth in Single-Cell Proteomics: A Cost-Effective Workflow and Expanded Framework for Data Evaluation"

Supplementary Table 1. Pearson correlation of protein intensity between bulk and single cell samples for each condition tested. For each comparison, the mean intensity across all single cell samples is calculated and compared to that of the bulk samples.

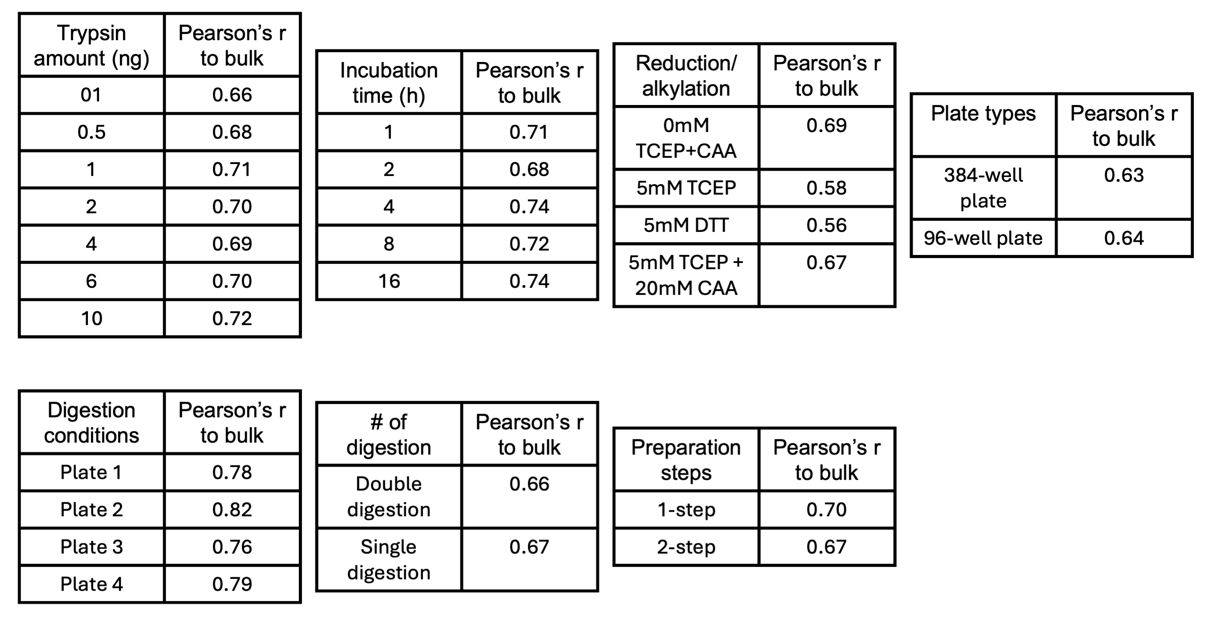
