## Supplementary Figure 14 for "Bridging Simplicity and Depth in Single-Cell Proteomics: A Cost-Effective Workflow and Expanded Framework for Data Evaluation"

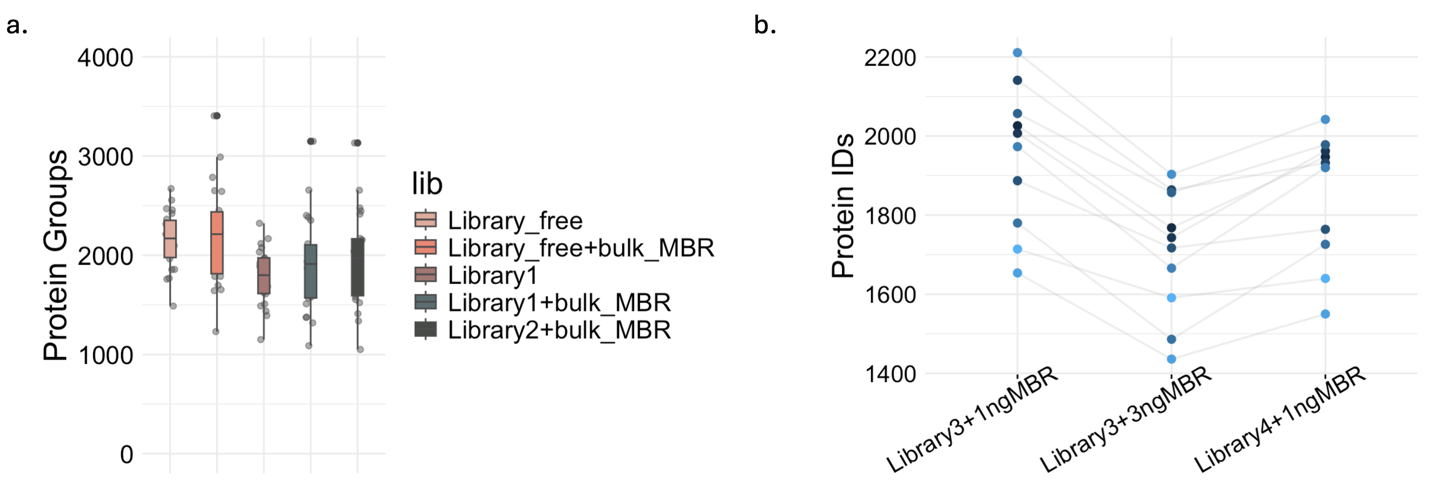

Supplementary Figure 14. Effect of DIA-NN search strategies. a, Average number of protein groups identified from single A549 cells acquired using a 15 min LC gradient (300 nL/min) with dia-PASEF (1.28 s cycle time), comparing different library settings with and without bulk Match-Between-Runs. Library 1: larger library generated using a 30 min LC method. Library 2: smaller, method-matched library generated using the same LC-MS/MS conditions as the single-cell runs. b, Comparison of three DIA-NN search strategies for 250pg HeLa injections. Library 3: larger library generated on timsTOF Ultra 2. Library 4: smaller, method-matched library generated using the same LC–MS/MS conditions as the 250pg HeLa runs.
