## Supplementary Figure 10 for "Bridging Simplicity and Depth in Single-Cell Proteomics: A Cost-Effective Workflow and Expanded Framework for Data Evaluation"

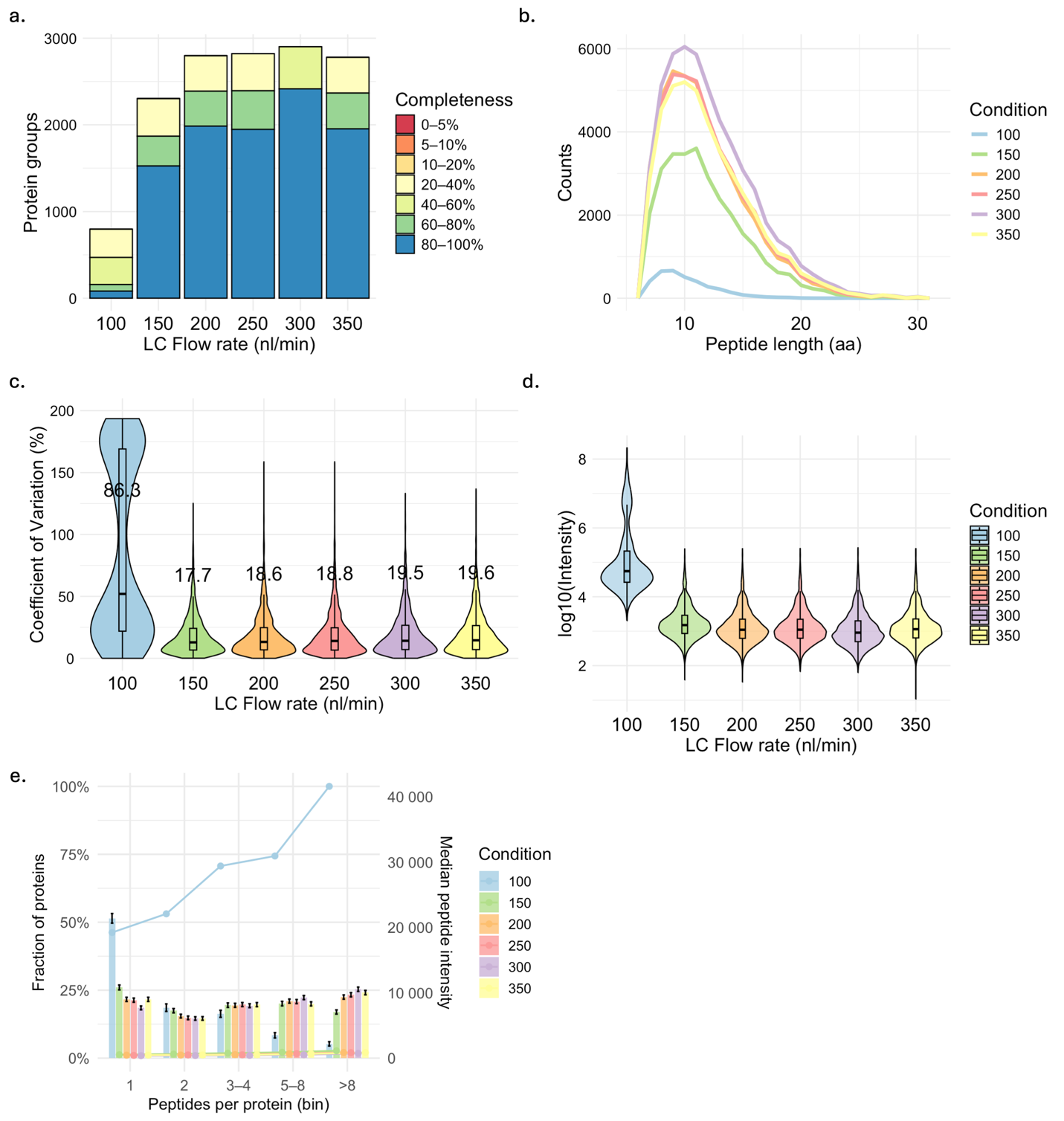

Supplementary Figure 10. Effect of LC flow rate (15min gradient) coupled with dia-PASEF window MS with 1.28s cycle time (250pg Hela per injection). a. Protein coverage completeness across cells. Stacked bars show the number of protein groups binned by completeness (0–5% to 80–100%); b. Peptide length distributions (PSM versus amino-acid length) across workflows; c. Coefficient of variation (CV) of protein intensities across injections shown with violin plots with median labeled for both conditions; d. Dynamic range of protein intensity between conditions; e. Peptide-per-protein ratio depth and median peptide intensity by conditions.
