## Supplementary Figure 7 for "Bridging Simplicity and Depth in Single-Cell Proteomics: A Cost-Effective Workflow and Expanded Framework for Data Evaluation"

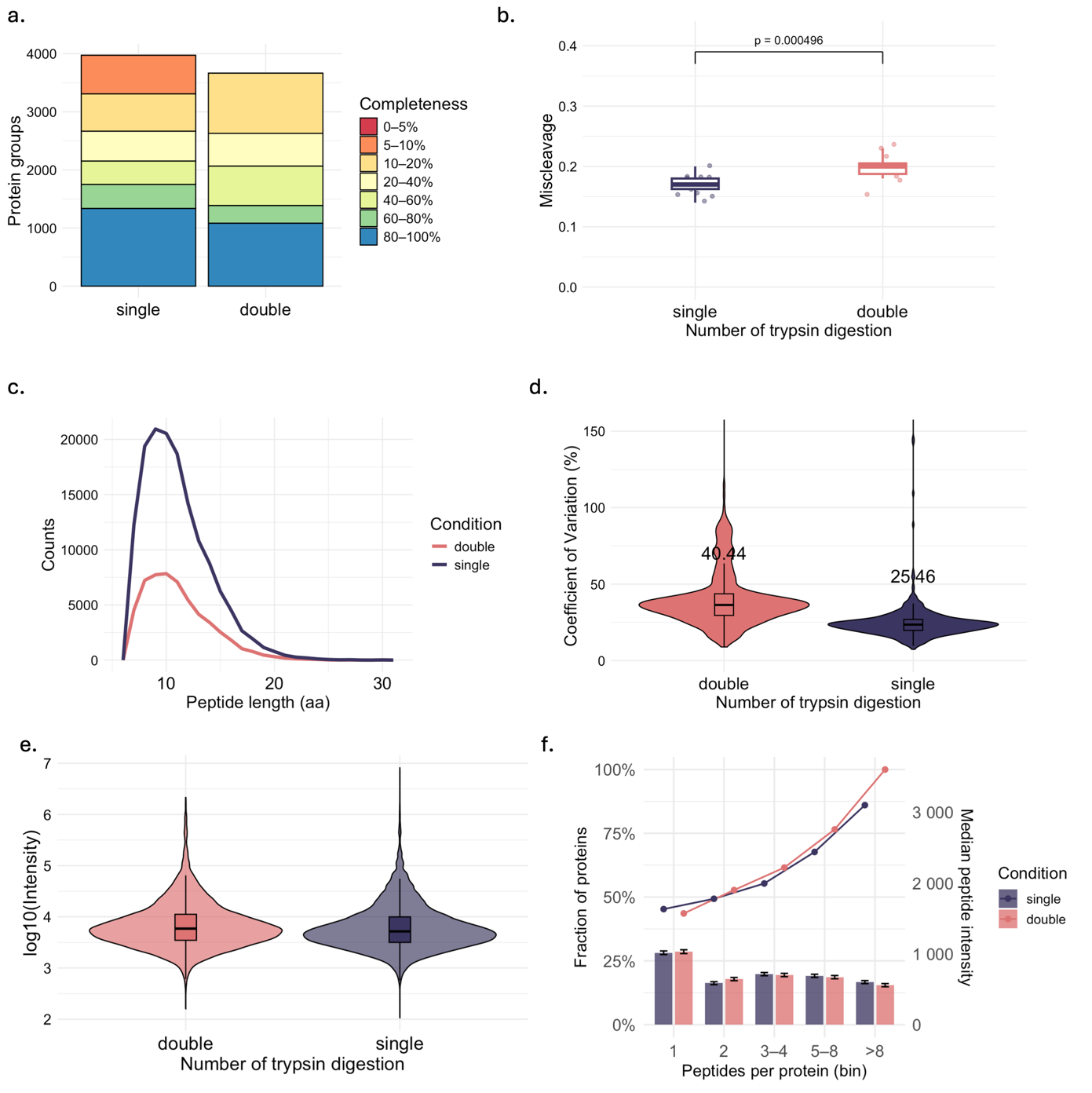

Supplementary Figure 7. Effect of single versus double trypsin digestion on single-cell proteomics data quality. a. Protein coverage completeness across cells. Stacked bars show the number of protein groups binned by completeness (0–5% to 80–100%); b. Tryptic missed-cleavage comparing single versus double trypsin digestion; c. Peptide length distributions (PSM versus amino-acid length) across workflows; d. Coefficient of variation (CV) of protein intensities across cells shown with violin plots with median labeled for each condition; e. Dynamic range of protein intensity between conditions; f. Peptide-per-protein ratio depth and median peptide intensity by conditions.
