## Supplementary Figure 6 for "Bridging Simplicity and Depth in Single-Cell Proteomics: A Cost-Effective Workflow and Expanded Framework for Data Evaluation"

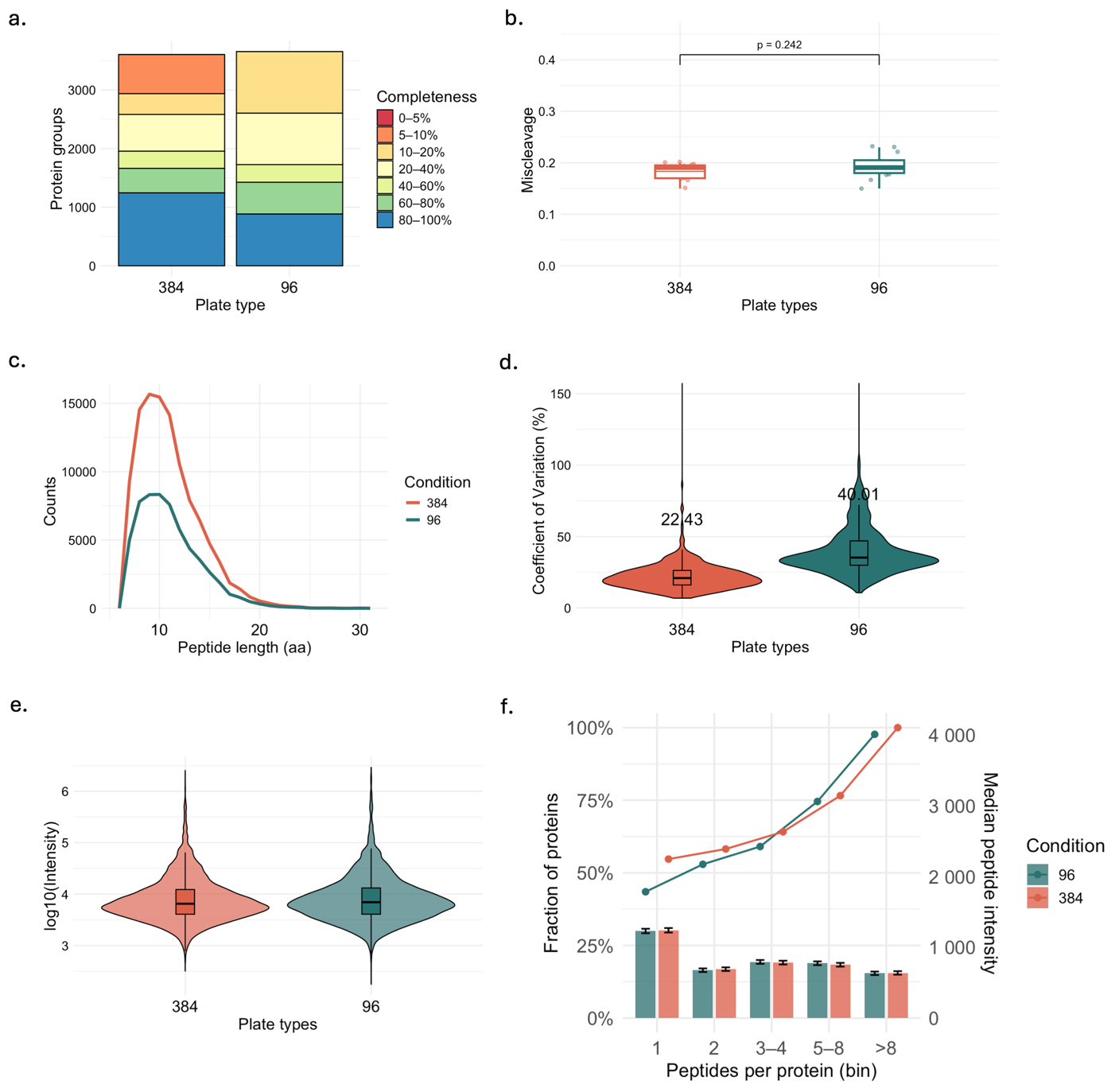

Supplementary Figure 6. Effect of plate type (384 versus 96-well plate) on single-cell proteome coverage and reproducibility. a. Protein coverage completeness across cells. Stacked bars show the number of protein groups binned by completeness (0–5% to 80–100%); b. Tryptic missed-cleavage of each plate type; c. Peptide length distributions (PSM versus amino-acid length) across different plate types; d. Coefficient of variation (CV) of protein intensities across cells shown with violin plots with median labeled for each condition; e. Dynamic range of protein intensity between conditions; f. Peptide-per-protein ratio depth and median peptide intensity by conditions.
