## Supplementary Figure 5 for "Bridging Simplicity and Depth in Single-Cell Proteomics: A Cost-Effective Workflow and Expanded Framework for Data Evaluation"

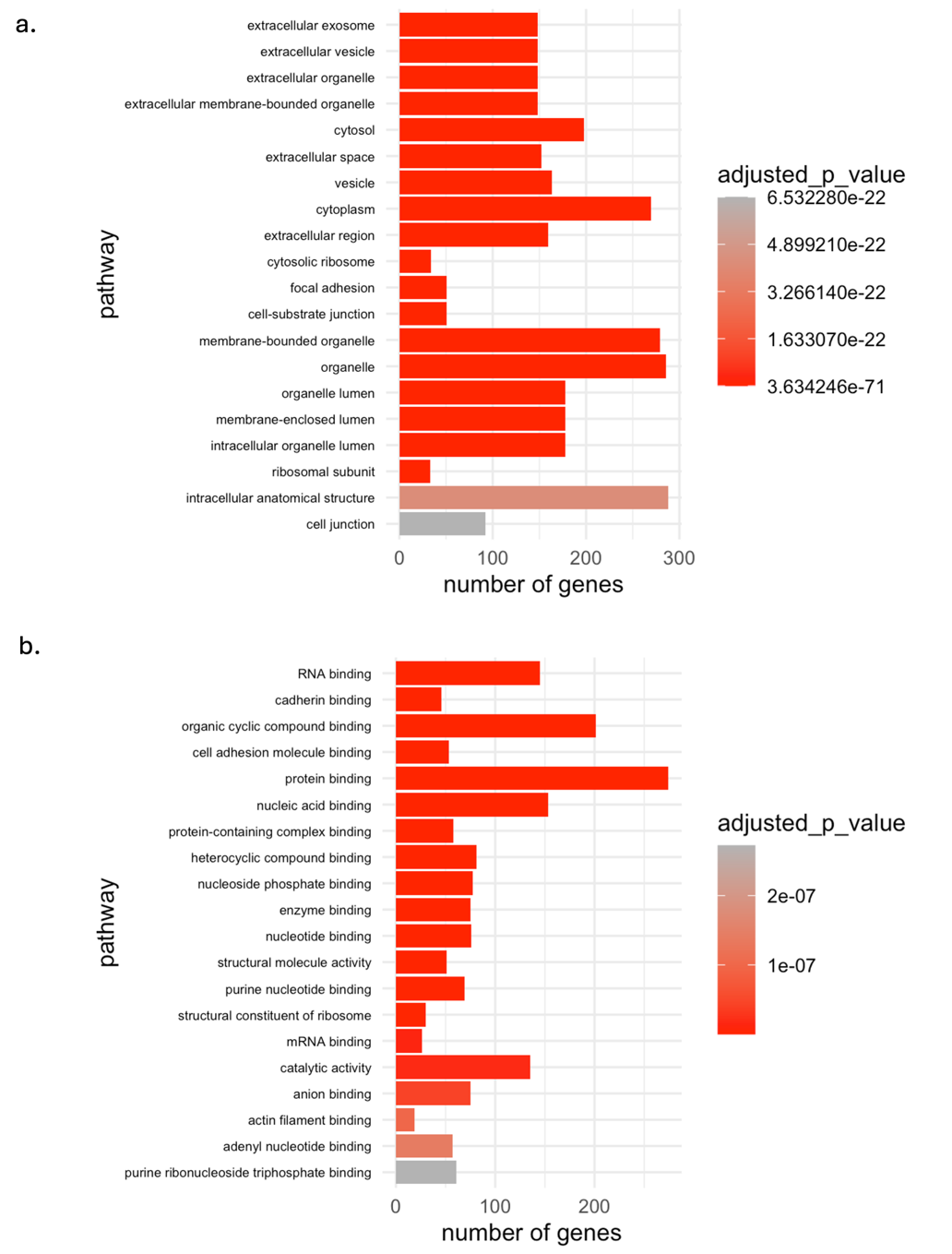

Supplementary Figure 5. GO analysis of proteins identified from reduction and alkylation. a. Cellular component GO analysis; b. Molecular function GO analysis.
