## Supplementary Figure 4 for "Bridging Simplicity and Depth in Single-Cell Proteomics: A Cost-Effective Workflow and Expanded Framework for Data Evaluation"

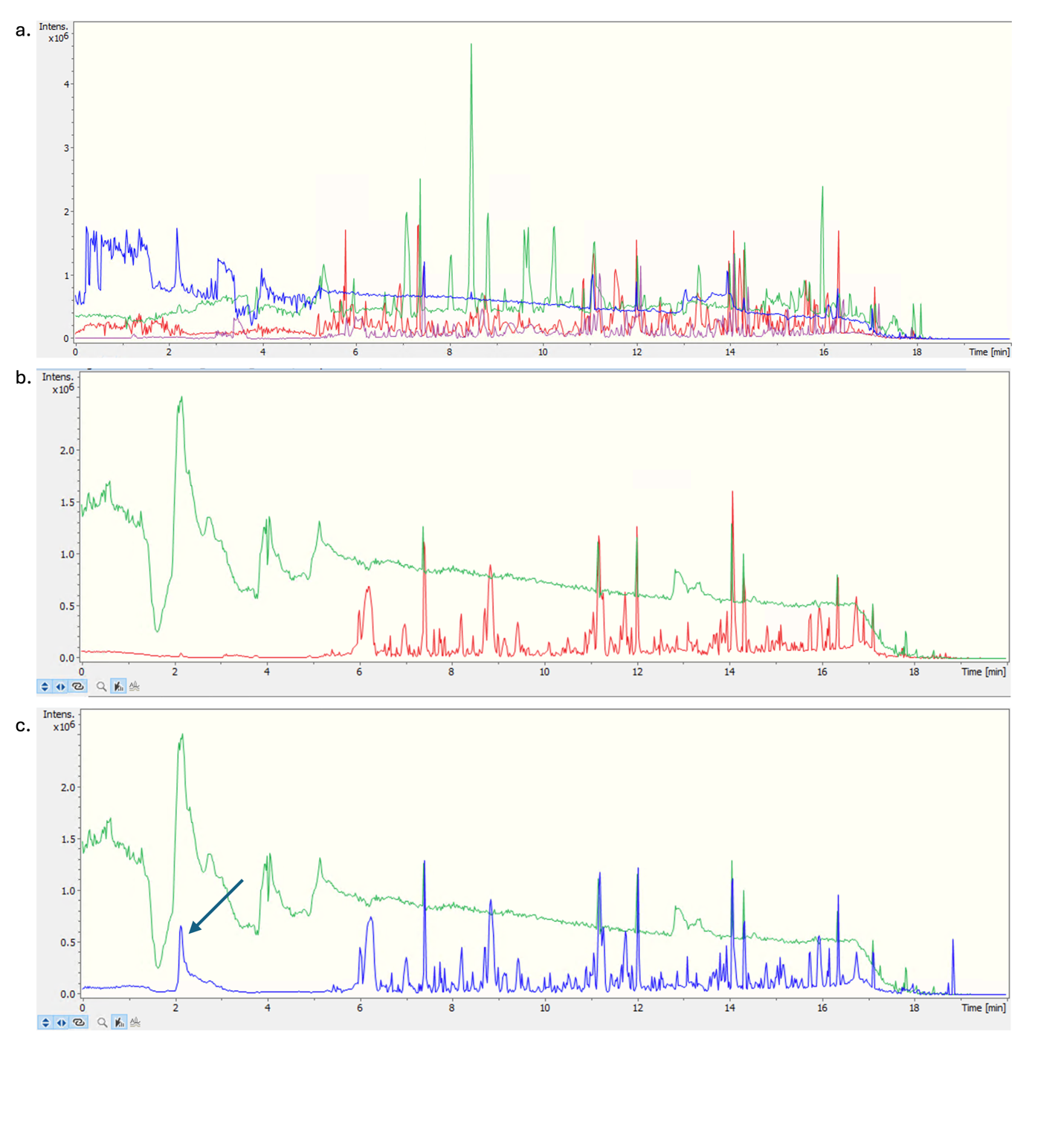

Supplementary Figure 4. Chromatograms for reduction and alkylation condition tests. a. An overlap view of the chromatogram for 0mMTCEP+CAA (purple), 5mM TCEP+20mM CAA (blue), 5mM TCEP (green) and 5mM DTT (red); b-c. Chromatograms showing contamination carryover. Samples with reduction and alkylation of 5mM TCEP+20mM CAA shown in green; sample injection before running reduced and alkylated samples (red, panel b) compared to one after running reduced and alkylated samples (blue, panel c).
