## Supplementary Table 3 for "Bridging Simplicity and Depth in Single-Cell Proteomics: A Cost-Effective Workflow and Expanded Framework for Data Evaluation"

Supplementary Table 3. Pearson correlation r values and R^2^ values for each condition plotted in Suppl. Figure 3.

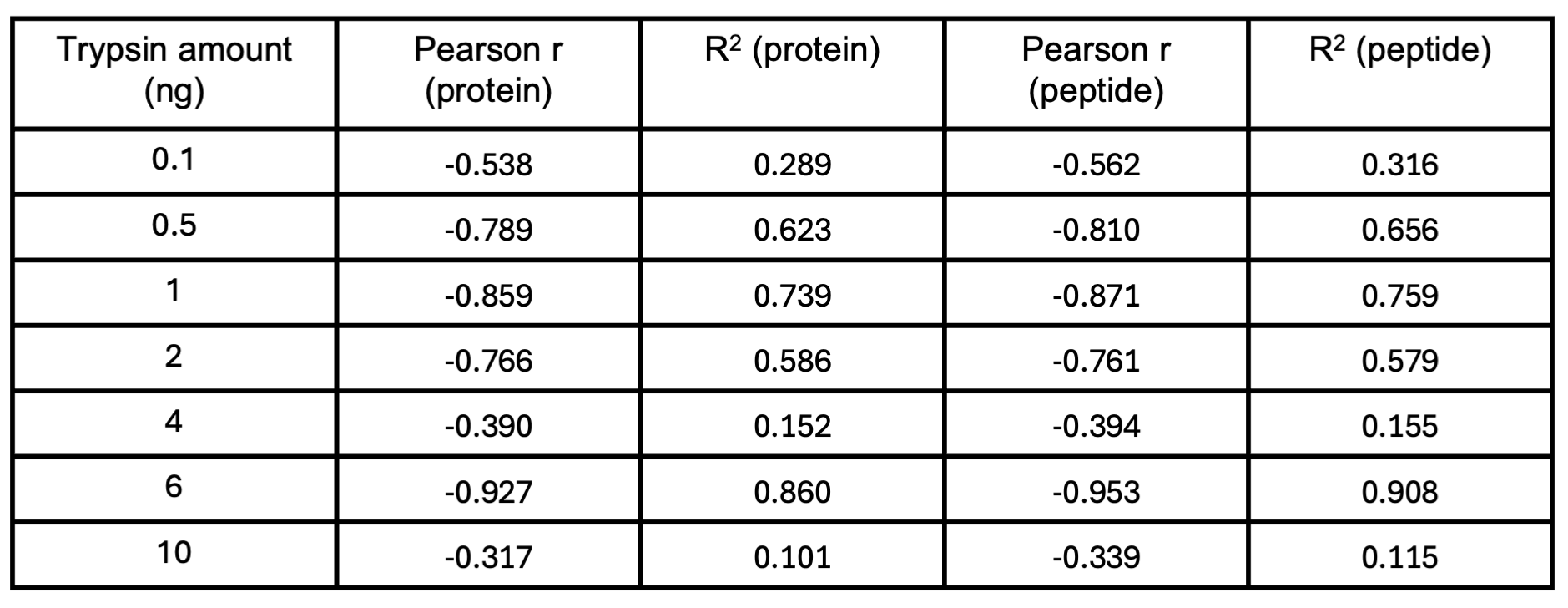
