## Supplementary Figure 3 for "Bridging Simplicity and Depth in Single-Cell Proteomics: A Cost-Effective Workflow and Expanded Framework for Data Evaluation"

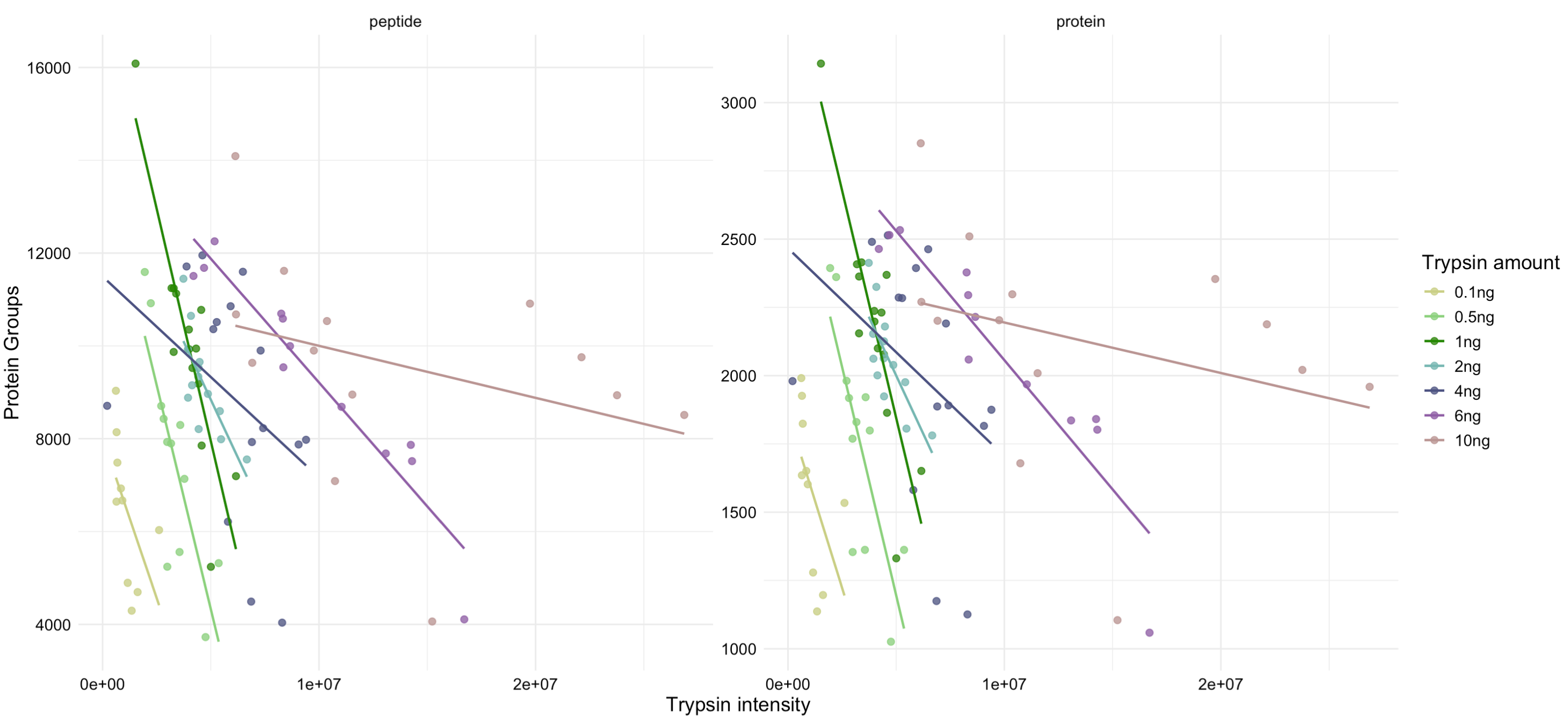
Supplementary Figure 3. Relationship between trypsin intensity and the number of protein IDs (right) and peptide IDs (left) for each condition.
