## Supplementary Figure 2 for "Bridging Simplicity and Depth in Single-Cell Proteomics: A Cost-Effective Workflow and Expanded Framework for Data Evaluation"

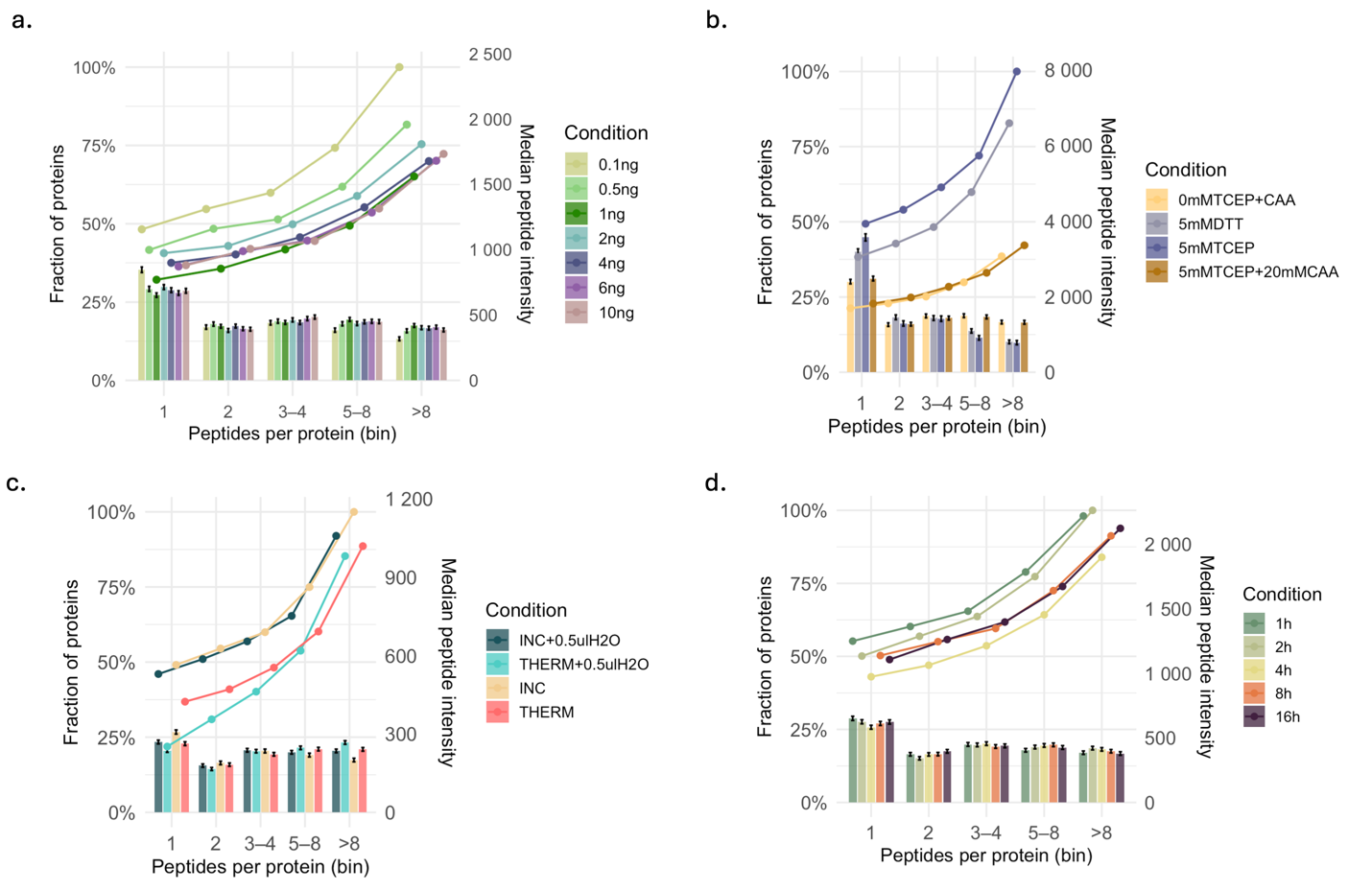

Supplementary Figure 2. Peptide-per-protein ratio depth and median peptide intensity by condition. Bar plots show the average number of peptides identified per protein across conditions and the overlaid dots indicate the corresponding median peptide intensity.
