## Supplementary Figure 1 for "Bridging Simplicity and Depth in Single-Cell Proteomics: A Cost-Effective Workflow and Expanded Framework for Data Evaluation"

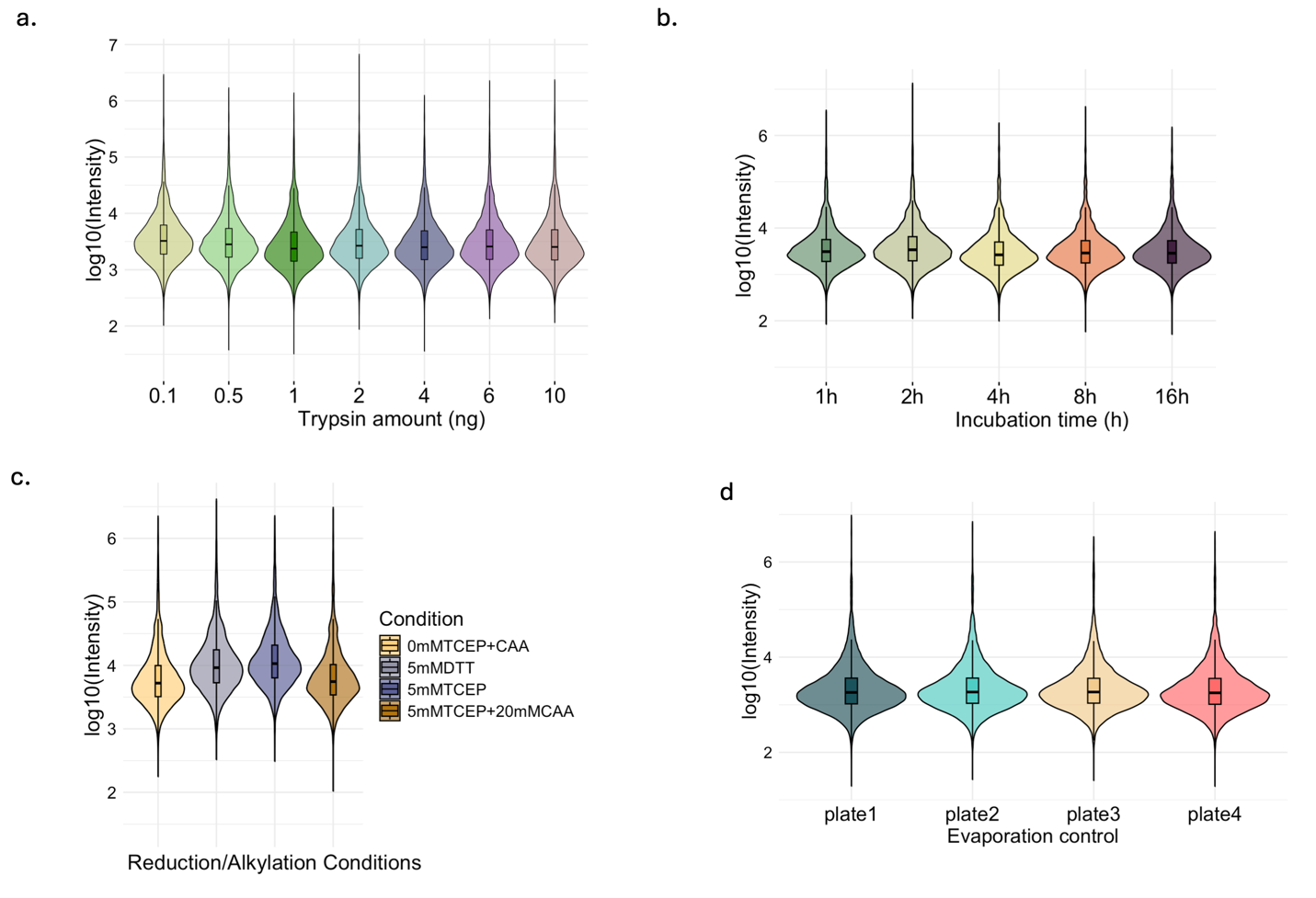

Supplementary Figure 1. Dynamic range of protein intensity across sample preparation parameters. Log_10_ transformed protein intensity distributions presenting dynamic range across different sample preparation conditions. Each distribution indicates the intensity spread and the central tendency, enabling comparison of overall protein trend and dynamic range across workflows.
